# *Helicobacter pylori* γ-glutamyltransferase relates to proteomic adaptions important for gastric colonization

**DOI:** 10.1101/2024.03.11.584369

**Authors:** Sonja Fuchs, Michaela K. Fiedler, Nicole Heiduk, Andreas Wanisch, Aleksandra W. Debowski, Barry J. Marshall, Stephan A. Sieber, Markus Gerhard, Raquel Mejías-Luque

## Abstract

*Helicobacter pylori* γ-glutamyltransferase (gGT) is a virulence factor that promotes bacterial colonization and immune tolerance. Although some studies addressed potential functional mechanisms, the supportive role of gGT for *in-vivo* colonization remains unclear. Additionally, it is unknown how different gGT expression levels may lead to compensatory mechanisms ensuring infection and persistence. Hence, it is crucial to unravel the *in-vivo* function of gGT. We assessed acid survival under conditions mimicking the human gastric fluid and elevated the pH in the murine stomach prior to *H. pylori* infection to link gGT-mediated acid resistance to colonization. By comparing proteomes of gGT-proficient and -deficient isolates before and after infecting mice, we investigated proteomic adaptations of gGT-deficient bacteria during infection. Our data indicate that gGT is crucial to sustain urease activity in acidic environments, thereby supporting survival and successful colonization. Absence of gGT triggers expression of proteins involved in the nitrogen and iron metabolism and boosts the expression of adhesins and flagellar proteins during infection, resulting in increased motility and adhesion capacity. In summary, gGT-dependent mechanisms confer a growth advantage to the bacterium in the gastric environment, which renders gGT a valuable target for the development of new treatments against *H. pylori* infection.

**Author Summary:** *H. pylori* γ-glutamyltransferase (gGT) is a virulence factor that strongly supports bacterial colonization. Despite considerable research on the function of gGT, the exact role of this enzyme in ensuring *in-vivo* infection remained elusive. We developed a novel system that allowed us to selectively inhibit gGT-activity and used this model to assess the function of gGT in the gastric environment. We found that gGT sustains urease activity in acidic environments thereby facilitating survival and effective colonization. In addition, we identified several compensatory mechanisms triggered by the loss of gGT which ensure colonization and persistence. These mechanisms included increased flagellar motility, adhesion capacity and expression of proteins involved in the nitrogen and iron metabolism. These findings unraveled novel functional roles of gGT important for bacterial colonization and thereby confirmed gGT as a promising target for novel treatments against *H. pylori* infection. By comprehensively addressing the compensatory mechanisms resulting from the loss of gGT-activity, the success of such new treatments can be improved.

## Introduction

*Helicobacter pylori* inhabits the stomach of more than 40% of the world’s population with varying prevalence across different regions [1]. Although most patients are asymptomatic, *H. pylori* causes gastric inflammation, elevating the risk for severe gastroduodenal diseases including ulcers, gastric cancer, or MALT lymphoma [1]. The severity of *H. pylori*-induced diseases is attributed to an array of virulence factors, including γ-glutamyl transferase (gGT) that was shown to contribute to gastric cancer development [2] and peptic ulcer disease [3].

*H. pylori* gGT catalyzes the hydrolysis of glutamine and glutathione, thereby releasing the metabolites glutamate, ammonia and cysteinyl glycine (Cys-Gly) into the bacterial microenvironment [4]. It is well-established that gGT-dependent alterations in metabolite concentrations affect host cells. Elevated ammonia levels exert cytotoxic effects on gastric epithelial cells [5, 6], while the release of reactive oxygen species resulting from gGT-activity triggers the secretion of pro-inflammatory interleukin 8 (IL-8) and the induction of apoptosis [3] in gastric epithelial cells. In addition to its effect on gastric epithelial cells, gGT-activity also facilitates immune evasion of the bacterium [7]*. H. pylori* gGT-induced glutamine deprivation hampers proliferation of T cells [8] and their effector function [9]. Simultaneously, a gGT-mediated increase in glutamate levels skews dendritic cells towards a more tolerogenic phenotype, favoring the proliferation of regulatory T cells that subsequently suppress effector T cells [10].

Beyond its impact on host cells, *H. pylori* gGT also promotes bacterial colonization, as evidenced by several studies demonstrating that *ggt* deletion diminishes colonization in murine infection models [5, 8, 11, 12]. However, even though *H. pylori* gGT is recognized as a key colonization factor, a small fraction of gGT-deficient bacteria is able to colonize and persist in the stomach at similar levels compared to the wild type strain (wt), indicating that the gGT-deficient *H. pylori* strain is somehow able to compensate for the loss of gGT-activity [8]. However, the underlying mechanisms that allow gGT-deficient bacteria to adapt in the absence of the enzyme, and to provide an advantage to the bacterium during colonization, remain elusive. Nevertheless, the fact that all gastric but not all enterohepatic *Helicobacter* express gGT suggests that the enzyme plays a central role specifically in the stomach environment [13].

A differential immune cell infiltration triggered by the gGT enzyme was proposed as a possible explanation for the initial colonization hurdle observed for gGT-deficient strains. However, this hypothesis was dismissed because a reduced colonization capacity of gGT-deficient bacteria was also observed in the absence of immune cells [8]. *In-vitro* studies further suggested that gGT might be metabolically important for the bacterium, providing another plausible explanation for the supporting effect of gGT during colonization. In this context, *H. pylori* gGT was shown to enhance bacterial acid survival *in-vitro* in the presence of urea [14] and to contribute to *H. pylori*’s carbon and nitrogen metabolism by enabling the bacterium to use extracellular glutamine. *H. pylori* is unable to take up glutamine directly but has to hydrolyze the amino acid to glutamate first, which is then taken up via the sole glutamate transporter GltS, and subsequently incorporated into metabolic pathways [4, 13]. The inability of *gltS*-deficient *H. pylori* to infect Mongolian gerbils stresses that this gGT-linked uptake system is also relevant *in-vivo*. Apart from glutamine, gGT-derived Cys-Gly might also support *H. pylori*’s metabolism as this metabolite has recently been shown to be internalized by the bacterium in a gGT-dependent manner [15] and has already been found to be essential for the proliferation of other pathogens during infection including *Francisella tularensis* [16] and *Neisseria meningitidis* [17].

Considering that different enzymatic activity levels have been observed in clinical isolates and that gGT activity has a high impact on bacterial growth and colonization, it is essential to identify compensatory mechanisms triggered in the absence of gGT, and to understand the metabolic role of this virulence factor in more detail. To this end, we have analyzed changes in bacterial protein expression occurring during infection of mice with wild type or gGT-deficient *H. pylori* and linked the contribution of gGT to acid resistance to colonization. Our findings indicate that gGT favors acid survival by supporting the key acid survival factor urease and has essential roles in different metabolic pathways important for the growth and infection capacity of *H. pylori*.

## Results

### Generation of gGT-inducible *H. pylori* strains

The analysis of the function of the virulence factor gGT has mainly relied on comparing *H. pylori* wild type strains (wt) with gGT-deficient deletion mutants (Δ*ggt*). However, this approach does not accurately reflect human infections, in which all clinical isolates were found to express gGT [13], albeit at varying levels, and the activity of the enzyme determines pathology [2, 3]. Therefore, a new model allowing for regulation of the expression levels of gGT was established to study the virulence factor gGT in more depth.

*H. pylori* strains with conditional gGT-activity were generated using the tetracycline-inducible gene expression system developed by Debowski et al. [18–20]. First, the *ggt* gene was placed under the control of the *tet*-inducible promoter *tetO1* (P_tetO1_), which is derived from the *H. pylori* urease promoter (P_ureAB_) but devoid of its transcriptional regulatory elements and contains the *tet* operator sequence (*tetO)* enabling binding of the *tet* repressor (*tetR*) [18]. To exchange the *ggt* promoter plasmids pGGT_rpsL-cat and pGGT_tetO1 were generated (**S1 FigS1 Fig**) and introduced into streptomycin-resistant *H. pylori* strains. Subsequently, the introduction of *tetR* into the *H. pylori* genome close to the *mdaB* locus effectively suppressed gGT gene expression (**Fig 1 A, Fig 1 B**). For *tetR* expression, the *flaA* (P_FlaA_) promoter was chosen because Debowski et al. found that *H. pylori* strains with a P_FlaA_-controlled *tetR* expression strongly expressed the target gene in the presence of an inducer, quickly reached maximum expression levels and responded to minimal inducer concentrations [18]. Correct insertion of *tetR* and *tetO1* into the genome resulting in *H. pylori* strain *g::O1 t* was confirmed by PCR (**S2 Fig**). The functionality of the system was assessed by measuring gGT-activity and protein expression in the presence and absence of the inducer anhydrotetracycline (ATc). *TetO1* mediated expression of gGT was determined to be 1.5 to 2-fold higher than in the corresponding wt strains (**Fig 1 B**). Without ATc no protein was detected in Western blots (**S3A Fig**) and enzymatic gGT activity was found to be below the detection limit (**Fig 1 B**). Replacement of P_ggt_ with P_tetO1_ without the introduction of *tetR* resulted in the strain *H. pylori g::O1 C* with constitutively high gGT-activity independent from the presence of an inducer, that was used as a control. Inducible strains expressed gGT even in the presence of ATc concentrations as low as 5 ng/ml (**Fig 1 C**). The strain G27 reached maximum gGT expression in the presence of 5 ng/ml of ATc, whereas 25 ng/ml were needed to fully induce gGT expression in PMSS1, and lower ATc concentrations only partially induced gGT-activity in this strain. As ATc is derived from the antibiotic tetracycline, it potentially has toxic effects on bacteria even in low concentrations. To test this, growth of *H. pylori* wt strains in the presence of different ATc concentrations was monitored. A concentration of 100 ng/ml did not influence bacterial growth compared to the solvent control, while higher amounts of ATc limited growth in a dose-dependent manner (**S3B Fig**). Hence, we selected a concentration of 100 ng/ml for further experiments, as this concentration 1) did not impact *H. pylori* growth, 2) fully induced gGT-activity, and 3) was determined by Debowski et al. to be necessary for maximum induction of *tetO1*-controlled GFP expression in *H. pylori* [18]. In the strain PMSS1 *g::O1 t* maximum induction of gGT-activity was achieved after 18 h in the presence of 100 ng/ml ATc, while a significant silencing of *ggt* gene expression after removal of the inducer was already observed after 8 h (**Fig 1 D**). In contrast to PMSS1, induction and silencing of gGT in the strain G27 *g::O1 t* was achieved already after 6 h (**S3C Fig**). In summary, these results demonstrate that functional *H. pylori* tet-on gGT strains and functional high gGT-expressing strains were generated.

**Fig 1.**
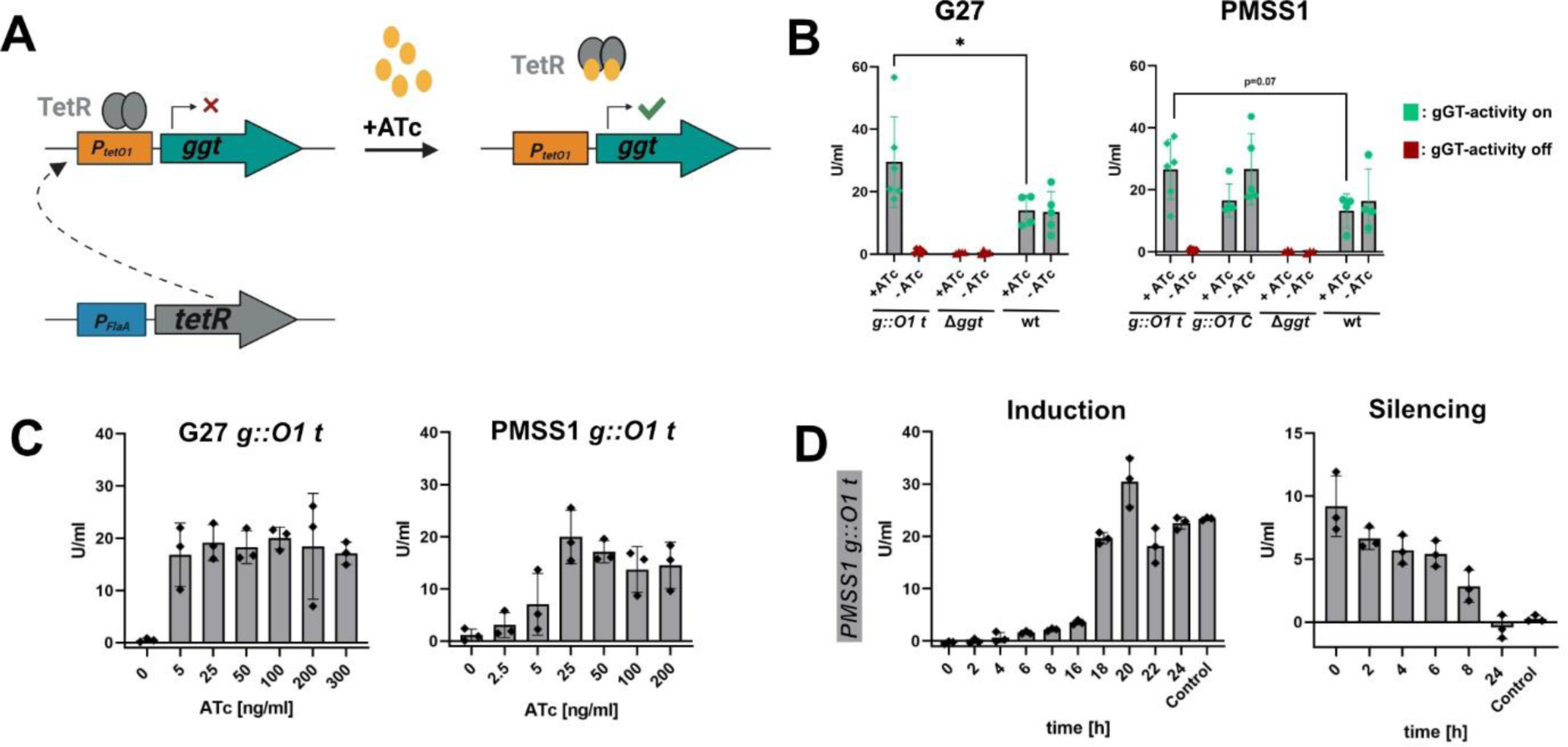
Establishment and characterization of inducible *tet*-on gGT *H. pylori* strains. (**A**) Graphic representation of the *tet*-on gGT system. The *ggt* promoter was replaced by the *tet* inducible promoter *tetO1* (P_tetO1_). TetR was introduced in the *H. pylori* genome and is controlled by a *flaA* promoter (P_FlaA_). TetR binds to *tetO1* blocking expression of gGT. The inducer anhydrotetracycline (ATc) binds to *tetR* resulting in disassociation of the repressor and activation of *ggt* expression. (**B**) gGT-activity of conditional gGT mutants. Bacteria were cultured in BHI/10% FCS in the presence of 100 ng/ml ATc for 24 h. *H. pylori g::O1 t* has *tetO1* and *tetR* integrated into its genome. In *H. pylori g::O1* C only the *ggt* promoter was replaced by *tetO1,* n=3-6. (**C**) gGT-activity in the presence of different ATc concentrations after 24 h of culture in BHI/10% FCS. The same amount of solvent (ethanol) was added to all cultures as a control, n= 3. (**D**) TetR-controlled induction and silencing of gGT-activity in PMSS1 *g::O1 t*. For induction, 100 ng/ml ATc was added to the culture (starting concentration: 1×10^8^ bacteria/ml). As a control, gGT-activity of PMSS1 *g::O1 t* after induction for 24 h was measured. For silencing, gGT-activity of PMSS1 *g::O1 t* was first induced for 24 h with 100 ng/ml ATc during growth in BHI/10% FCS. The culture was washed with PBS and resuspended in ATc-free medium at a concentration of 1*10^8^ bacteria/ml. As a control, an uninduced 24 h old culture of PMSS1 *g::O1 t* was used, n=3. Bars represent the mean of three to six independent experiments as represented by single dots. Error bars represent the standard deviation. n: number of independent experiments. Mann-Whitney U-test. *p<0.05.

### *H. pylori* gGT is important for initial colonization and for persistence

*H. pylori* gGT is a virulence factor that facilitates initial colonization [8, 11, 12]. However, Wüstner et al. observed similar infection levels of a wt and a gGT-deficient strain in a mouse model after 1 month of infection, despite an initial colonization hurdle of the gGT-deficient strain [8]. This raised the question during which infection phases gGT is important for *H. pylori* colonization. To address this question, mice were infected for up to 6 months either with a wt-, a gGT-deletion strain or a strain expressing high levels of gGT (*g::O1 C*) (**Fig 2**). At the beginning of the infection, colonization was significantly lower in mice infected with the gGT-deletion strain compared to those infected with PMSS1 wt and PMSS1 *g::O1 C*, confirming previous findings about the role of gGT during initial infection. However, colonization levels of the gGT-deletion strain increased over time, peaking after one month of infection, consistent with observations by Wüstner et al. Subsequently, after this colonization peak, infection levels in mice infected with PMSS1 Δ*ggt* decreased indicating that gGT also contributes to persistence. After 3 months and 6 months of infection colonization levels of the gGT-deficient strain were significantly lower than those of the wt and gGT-high strain. No difference in colonization was observed comparing the infection levels of mice inoculated with the wt or high gGT-expressing strain throughout the course of the infection.

**Fig 2.**
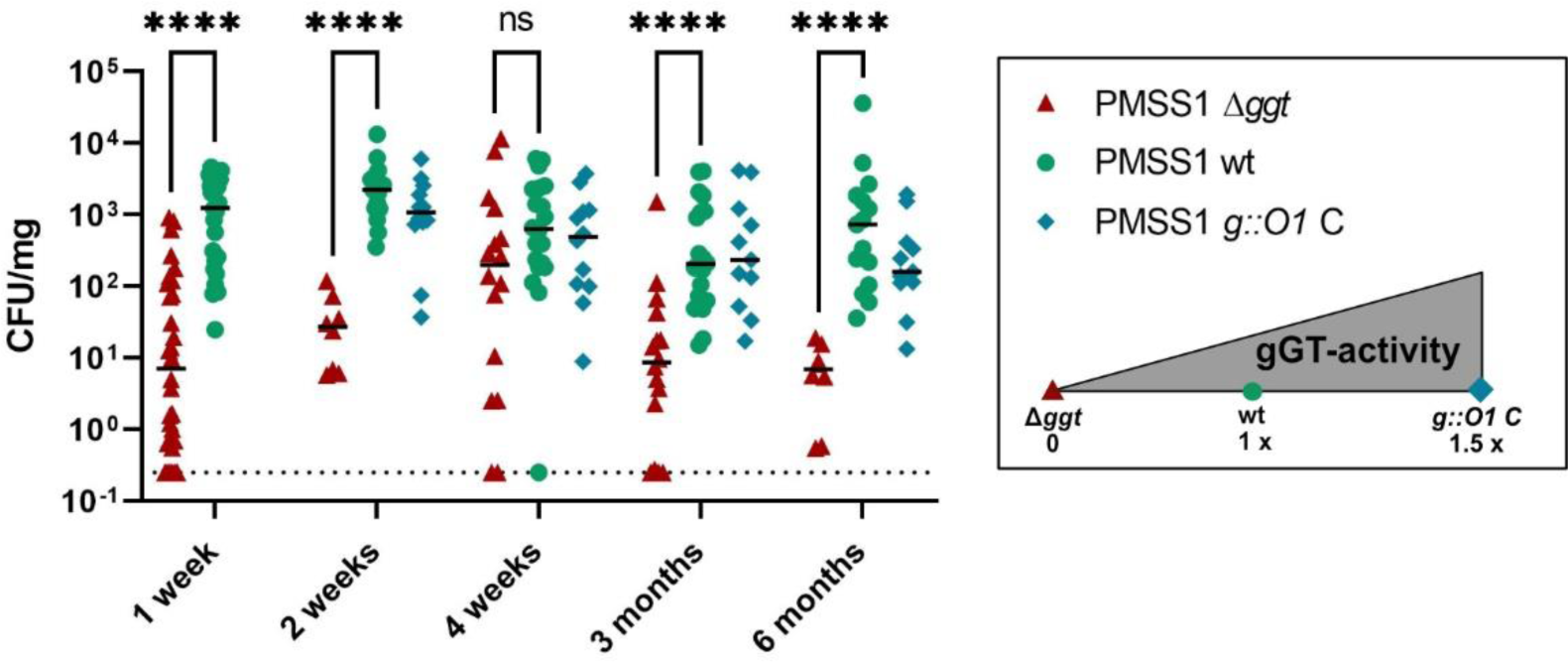
*H. pylori* gGT contributes to initial colonization and persistence. Colony forming units (CFU) in the stomach (mg) of C57BL/6 mice after infection with PMSS1 wt, Δ*ggt* or *g::O1 C.* Mice were inoculated with 2*10^8^ bacteria twice and were sacrificed after a defined amount of time (1 week - 6 months) after the first infection. Colonization was assessed by plating stomach homogenates. The detection limit is indicated by a dotted line. Each dot represents one individual mouse. PMSS1 wt (n=15-27), PMSS1 Δ*ggt* (n=8-32), PMSS1 *g::O1 C* (n=12), n: number of mice. Mann-Whitney U-test. ****p<0.01.

In summary, these findings suggest that gGT is not only crucial for initial colonization but also for persistence.

### *H. pylori* gGT contributes to acid resistance supporting *in-vivo* colonization

Next, we hypothesized, based on the study by Miller and Maier, that gGT is important for acid resistance which might contribute to the effect of this virulence factor on initial colonization [14]. *H. pylori* gGT was described to enhance the survival of strain 26695 at pH 3 in the presence of high urea concentrations [14]. However, this study does not fully account for the complexity of the human stomach, where the gastric fluid pH ranges from 1.7 to 6 depending on food intake [21], and additional components of the gastric fluid such as the proteolytic enzyme pepsin [22] or the bile acid sodium taurocholate [23] further challenge *H. pylori* acid survival and growth. Consequently, simulation of stomach-like conditions is important to understand the role of gGT in acid survival. First, survival assays were performed with a physiologically relevant urea concentration of 8 mM [24, 25]. Under these conditions, both wildtype G27 and PMSS1 exhibited enhanced survival rates compared to isogenic gGT-deficient strains (**Fig 3 A**), confirming previous observations by Miller and Maier using a urea concentration of 20 mM [14]. PMSS1 wt demonstrated robust survival for up to 6 h, while the viability of the gGT-deficient mutant began to diminish after 2 h. The G27 strain was more vulnerable to acidity compared to PMSS1 under these conditions. Here, viable wt bacteria persisted even after 6 h, but no gGT-deficient mutants could be recovered after the same time interval (**Fig 3 A**). To confirm that McIlvaine buffer at pH 3 with 8 mM urea poses an acidic challenge to *H. pylori*, urease-deficient strains, which are well-known to be highly sensitive to acidic conditions [26], were used as internal controls, and died within an hour in this milieu as expected. Moreover, to rule out that gGT’s impact on acid resistance was due to a general survival defect of gGT-negative strains in McIlvaine buffer, survival was also monitored at pH 5 (**S5A Fig**). At this pH, the urease-deficient control strains and the gGT-deficient knock-out strains survived equally well as the wt for at least 4 h. To investigate whether the effect of gGT on acid resistance depends on the presence of urea, survival at pH 3 was assessed without supplements (**S5B Fig**). A supporting effect of gGT on *H. pylori’s* acid survival was evident but was less pronounced than in the presence of urea. After assessing the effect of gGT on acid survival with the laboratory strains PMSS1 and G27, the survival of clinical isolates with different gGT-activities was measured, revealing a significant correlation between gGT-activity and acid survival at pH 3 in McIlvaine buffer supplemented with physiologic concentrations of urea (**Fig 3 B**).

**Fig 3.**
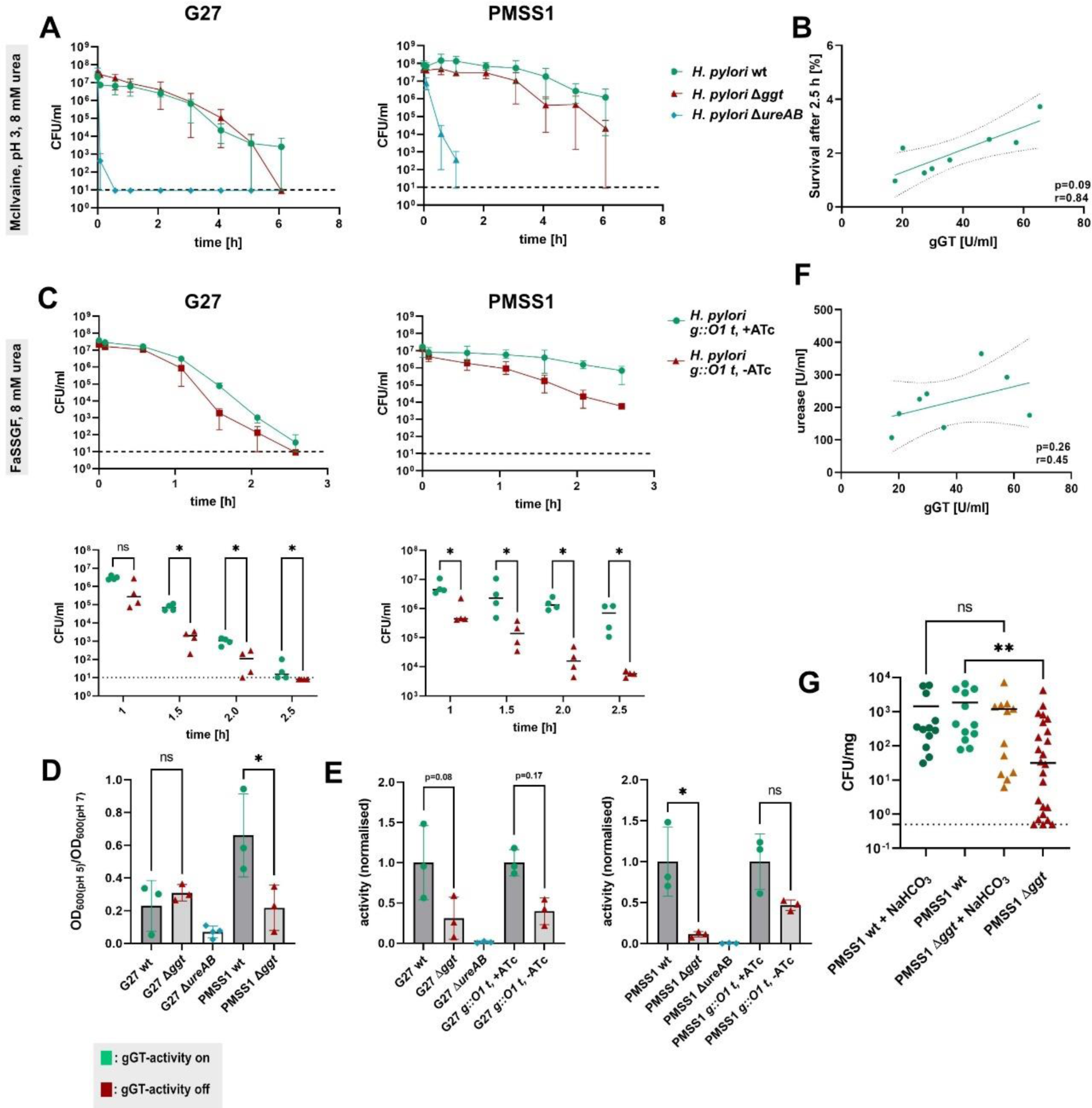
*H. pylori* gGT contributes to acid survival supporting stomach colonization. (**A**) Survival of *H. pylori* in McIlvaine buffer (pH 3) supplemented with 8 mM urea, n=3. The detection limit of the assay is indicated by a dotted line. (**B**) Correlation of survival of clinical isolates after 2.5 h with gGT-activity. Survival was determined in McIlvaine buffer (pH 3) supplemented with 8 mM urea. Each dot represents the mean survival of one clinical isolate, n=3. (**C**) Survival of *H. pylori* in fasted state simulated gastric fluid (FaSSGF) supplemented with 8 mM urea. The detection limit of the assay is indicated by a dotted line., n=4. (**D**) Growth of *H. pylori* in Brucella broth supplemented with 20% FCS at pH 5 normalized to the growth in Brucella broth at pH 7. Growth was quantified spectrophotometrically after 30 h. n=3. **(E**) Urease activity after exposure of *H. pylori* to pH 5. Strains were grown in BHI with 10% FCS, resuspended for 2 h in McIlvaine buffer (pH 5) and lysed for the detection of urease activity. Data was normalized to the mean activity of *H. pylori* wt or tet-on *H. pylori g::O1 t*. n=3. (**F**) Correlation of urease- and gGT-activity of clinical isolates. Each dot represents the mean activity of an isolate. n=3. (**G**) Colony forming units (CFU) in the stomach (mg) of C57BL/6 mice after infection with PMSS1 wt or Δ*ggt*. Mice were inoculated with 2*10^8^ bacteria and were sacrificed after one week of infection. Two groups of mice received sodium bicarbonate (NaHCO_3_) before the infection to elevate the pH in the stomach. Colonization was assessed by plating stomach homogenates. The detection limit is indicated by a dotted line. Each dot represents one individual mouse. n: number of independent experiments. Pearson’s correlation (B, F); Mann-Whitney U-test (C, G); ANOVA (D, E) * p<0.05, **p<0.01.

Next, we monitored survival under conditions similar to the gastric environment. For this purpose, fasted state simulated gastric fluid (FaSSGF) that includes constituents like pepsin, low concentrations of bile salts, and lecithin in physiological concentrations was used. FaSSGF has a comparable pH, osmolarity, and surface tension [27, 28] as the gastric fluid in the fasted state. G27 showed reduced survival rates in this medium compared to PMSS1, however, differences in survival between wt and gGT-deficient strains starting after 1.5 h showed that gGT-activity significantly supported survival of *H. pylori* strains PMSS1 and G27 in FaSSGF. (**Fig 3 C, S5C Fig**).

To infect the host *H. pylori* needs to survive the harsh environment of the stomach lumen for a short period. However, after arriving at the gastric mucosa, the bacterium is replicating in a rather neutral environment. Nevertheless, there is a pH gradient along the mucosa [24] and the mucosal pH can drop after food intake [29]. Thus, *H. pylori* has developed mechanisms enabling growth at mildly acidic conditions [30, 31]. To analyze the function of gGT in this context, we monitored *in-vitro* growth of *H. pylori* strains G27 and PMSS1 in the presence and absence of gGT at pH 5. BHI medium with 20% FCS did not allow for growth of *H. pylori* in these conditions (**S4 Fig**), but Brucella medium supplemented with 20% FCS supported growth of PMSS1 and G27. A low pH reduced growth of the wt strains (to 80% for PMSS1, to 20% for G27) compared to neutral conditions. *H. pylori* gGT strongly supported growth of PMSS1 while there was no observable effect of gGT on the strain G27 in the tested conditions. (**Fig 3 D**). In addition, growth of wt strains was accompanied by a strong increase in the pH of the medium while strains deficient of gGT elevated the medium pH to a lower extent (**S4 FigB Fig**).

Urease activity is a central mechanism of *H. pylori* enabling the survival [32] and growth [30] of the bacterium at low pH by catalyzing the hydrolysis of urea to ammonia and carbon dioxide. As gGT supports acid survival (**Fig 3 A-C**) and contributes to pH increase in the presence of urea (**S5D Fig**), the reduced survival in the absence of gGT-activity could be induced by impaired urease activity. To test this hypothesis, we evaluated urease activity after exposing *H. pylori* to pH 5 in McIlvaine buffer. Urease is considered to be a cytoplasmic protein, but 10-30% of the enzyme is also reported to be localized on the bacterial surface [33, 34]. Urease activity was notably reduced by 50% in *H. pylori* lysates (**Fig 3 E**) and in the supernatants (**S5E Fig**) in gGT deletion strains compared to wt strains.

Since both wt and gGT-deficient strains exhibited comparable survival at pH 5 in McIlvaine buffer (**S5A Fig**), variations in urease activity cannot be attributed to different survival modalities of wt and gGT-deficient strains. The reduction in urease activity was found in deletion strains and in the *H. pylori g::O1 t* model in the absence of ATc, suggesting a functional link between urease and gGT. This diminished urease activity was accompanied by a lower amount of this enzyme present in the supernatants of gGT-deficient *H. pylori* strains and, in the case of the strain G27, by a lower urease concentration in the lysate, as detected by Western blot (**S5F Fig**) and proteome analysis (**S6 Fig**). However, in the PMSS1 lysate, reduced urease activity could not be linked to decreased protein expression, suggesting other control mechanisms. To assess whether the activity level of gGT, in addition to its presence, is important for urease function, urease activity was assessed in clinical isolates. Urease activity tended to correlate with gGT-activity, even though this correlation was not found to be significant (**Fig 3 F**).

Taken together, our data revealed a protective function of gGT in acidic environments, supporting bacterial survival and growth. To link these findings to *in-vivo* colonization, we used a murine infection model in which C57BL/6 mice were administered sodium bicarbonate by oral gavage prior to infection to elevate the gastric pH. Following the infection of mice with wt or gGT-deficient PMSS1, colonization levels were assessed by plating stomach homogenates. While the wt strain did not benefit from sodium bicarbonate treatment, the infection level of the gGT-deficient strain notably increased after the treatment, overcoming the initial colonization hurdle of gGT-deficient PMSS1 (**Fig 3 G**).

Thus, our findings collectively indicate that gGT supports *H. pylori* colonization in acidic environments, partly by maintaining urease activity.

### Absence of *H. pylori* gGT leads to an upregulation of proteins involved in the nitrogen and iron metabolism

Even after elevation of the pH in the stomach, the colonization levels observed in mice infected with the gGT-deficient strain still tended to be lower than in mice infected with the wt strain (**Fig 3 G**), indicating that gGT might also support colonization by other mechanisms independent from acid resistance. To identify plausible additional functions of gGT or compensatory mechanisms allowing gGT deficient bacteria to colonize, isolates retrieved from the murine stomach were compared to the aliquots initially used for infection. The infection levels of mice used for this experiment followed the same trend as described in the previous section (**S7A Fig**).

To study protein expression profiles, re-isolates retrieved from different mice after 1 week, 1 month or 3 months of infection and the strains initially used for infection were grown until the early stationary phase (**S7B Fig**), lysed and analyzed by liquid chromatography–mass spectrometry (LC-MS/MS).

Alterations in the expression levels of various proteins implicated in the nitrogen metabolism were observed when comparing the proteome of PMSS1 wt and PMSS1 Δ*ggt*. *H. pylori* employs several enzymes to produce ammonia, which is subsequently assimilated by the bacterium [35]. This enzymatic ensemble encompasses gGT, formamidase (AmiF), amidase (AmiE), urease (UreAB), aspartate-ammonia lyase (AspA) and asparaginase (AsnA). Notably, gGT deficient strains showed significantly elevated expression levels of both AmiF and SdaA, and a discernible but not significant increase in the expression of AmiE (**Fig 4 A**). The altered expression of AmiE and SdaA was present pre-infection and remained to be similarly different between wt and gGT deficient strains after infection. However, AmiF expression diminished during the infection phase.

**Fig 4.**
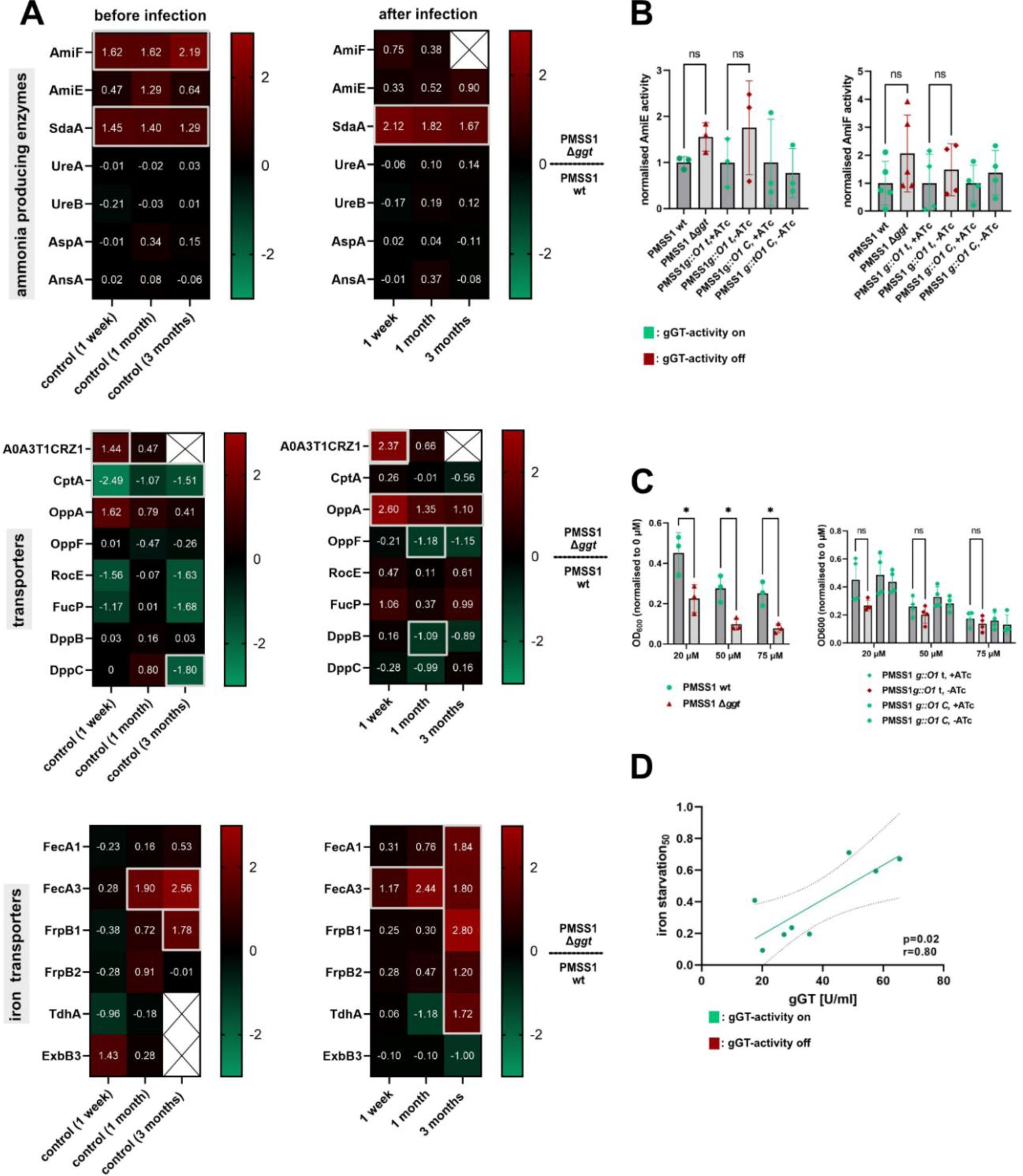
Proteins involved in the nitrogen and iron metabolism are differentially expressed in the absence of gGT. (**A**) Log_2_ transformed fold change of ammonium producing enzymes, peptide/metabolite transporters and of proteins involved in metal acquisition comparing the proteome of PMSS1 wt and Δ*ggt*. The proteomes of three single clones isolated after 1 week, 1 month and 3 months of infection from different mice were determined. Significant differences (-log_10_(p-value) > 1.3 and log_2_(fold change) ≥ I1I) are marked with grey boxes. Missing data points are colored white and are marked with a cross. (**B**) AmiE and AmiF activity of *H. pylori* normalized to the mean activity of *H. pylori* wt or tet-on *H. pylori g::O1 t.* (**C**) Growth of *H. pylori* in iron-restricted medium. Deferoxamine mesylate (DFO) was added to the culture in different concentration (20 µM, 50 µM and 75 µM). Growth was monitored spectrophotometrically and was normalized to the growth without supplementation. Bars represent the mean of three to four independent experiments as indicated by dots. Error bars represent the standard deviation. (**D**) Correlation of growth in iron-limited conditions with gGT-activity. Each dot represents the mean growth of one clinical isolate as measured in three independent experiments. Clinical isolates were grown in the presence of 50 µM DFO and growth data were normalized to the growth without supplementation. Students t-test (A; C). ANOVA (B); Pearson’s correlation (D). *\** p<0.05.

To confirm the changes in expression levels of ammonium producing enzymes detected by mass-spectrometry, AmiE and AmiF activities were measured in the presence and absence of gGT in the strain PMSS1. Both enzyme activities were increased 1.5 to 2-fold in the absence of gGT, when gGT was either deleted or its expression repressed by TetR (**Fig 4 B**). Moreover, enzymatic activity assays strengthened the finding that AmiF activity is reduced in re-isolates, while AmiE activity remained unchanged (**S8B Fig**). To understand whether altered AmiF and AmiE activities in the absence of gGT generally apply to *H. pylori* or whether this effect is strain-specific, activity measurements were also conducted with the G27 strain. Analogous to the observations in PMSS1, AmiF activity tended to be increased in the G27 strain in the absence of gGT. However, the effect of AmiE differed in G27, as the enzymatic activity was decreased upon silencing or deletion of gGT (**S8A Fig**).

In concert with ammonium producing enzymes, peptide transporters ensure the supply of the bacterium with nitrogen sources [35]. In this context, we detected differential expression of peptide transporters CptA and OppA in the gGT-deficient deletion mutant compared to the wt strain. CptA is known to transport the gGT-derived peptide Cys-Gly in *C. jejuni* [36], while OppA was predicted to be an oligopeptide transporter but its specificity in *H. pylori* remains unknown [37]. Prior to infection, CptA expression was very low in gGT-deletion mutants, but it reverted to wt levels during infection (**Fig 4 A**). Conversely, OppA, expression was found to be significantly higher in gGT-deficient compared to wt re-isolates subsequent to infection. In conclusion, the absence of gGT prompts *H. pylori* to orchestrate adjustments in expression profiles of various proteins implicated in nitrogen assimilation, indicating a central function of gGT in the nitrogen metabolism.

In addition to the alterations in the expression of proteins associated with the nitrogen metabolism, our experiment also revealed differential expression of iron-assimilating proteins in the absence of gGT. Among these is FecA3. The absence of gGT led to an upregulation of *fecA3* expression, which is a potential iron-citrate transporter [38], both pre- and post-infection (**Fig 4 A**). Furthermore, significantly increased expression of other proteins associated with iron acquisition, including the potential iron(III)citrate transporter FecA1, heme transporters FrpB1 and FrpB2 and the TonB-dependent heme receptor A (TdhA), was evident in gGT-deficient mutants after 3 months of infection (**Fig 4 A**).

These findings together with the prediction that gGT is controlled by the central iron regulator Fur [39], raised the question whether gGT plays a role in the bacterial iron metabolism. To address this question, we cultivated *H. pylori* strains under iron-restricted growth conditions with varying concentrations of the iron chelating agent deferoxamine mesylate supplemented to the growth medium. This experiment revealed that gGT-proficient strains grew significantly better under iron limiting conditions compared to their isogenic gGT-deficient mutant strains (**Fig 4 C**). Moreover, the gGT-activity of clinical *H. pylori* isolates correlated with the ability to grow in iron-limited conditions (**Fig 4 D**). The observed upregulation of iron acquisition proteins in gGT-deficient mutants might seem counterintuitive in this context. However, Pich et al. hypothesized that gGT-activity is involved in reducing Fe^3+^ to Fe^2+^ [38], which is subsequently assimilated by the FeoB transporter [40]. In this context, the elevated expression of specific proteins related to iron-acquisition could be a compensatory mechanism facilitating increased Fe^3+^ uptake. To validate whether gGT-deficient strains are more capable of taking up certain Fe(III) iron sources, we quantified the uptake of Fe(III)citrate into *H. pylori* and observed that gGT-deficient G27 and PMSS1 strains indeed display an 1.3-1.4x enhanced uptake of this iron source compared to their respective wt strains (**S8C Fig**). Taken together, our findings suggest that gGT confers a growth advantage in iron-deprived environments, which might be partially counterbalanced by an increased uptake of Fe(III) under iron rich conditions.

### Flagellar motility and adhesion capacity are enhanced in the absence of gGT subsequent to infection

In addition to its impact on the expression of proteins related to the bacterial metabolism, the absence of gGT also affected bacterial traits that have not been linked to gGT-activity yet, including flagellar proteins and adhesion factors.

Comparative analysis of the proteome of wt and gGT-deficient PMSS1 prior to infection revealed an upregulation of flagellar proteins in the wt strain (**Fig 5 A**). Mainly, proteins from the flagellar filament, including flagellin A (FlaA), B (FlaB) and G (FlaG), and from the flagellar hook, including FlaG, were found to be affected by the absence of gGT. Increased secretion of the same flagellar proteins was observed for *H. pylori* Δ*ggt* G27 and PMSS1 strains as compared to the wt after exposure to pH 5 (**S6 Fig**), implying that the observed increased presence of flagellar proteins in the cytoplasm of the wt strains may result from a difference in secretion.

**Fig 5.**
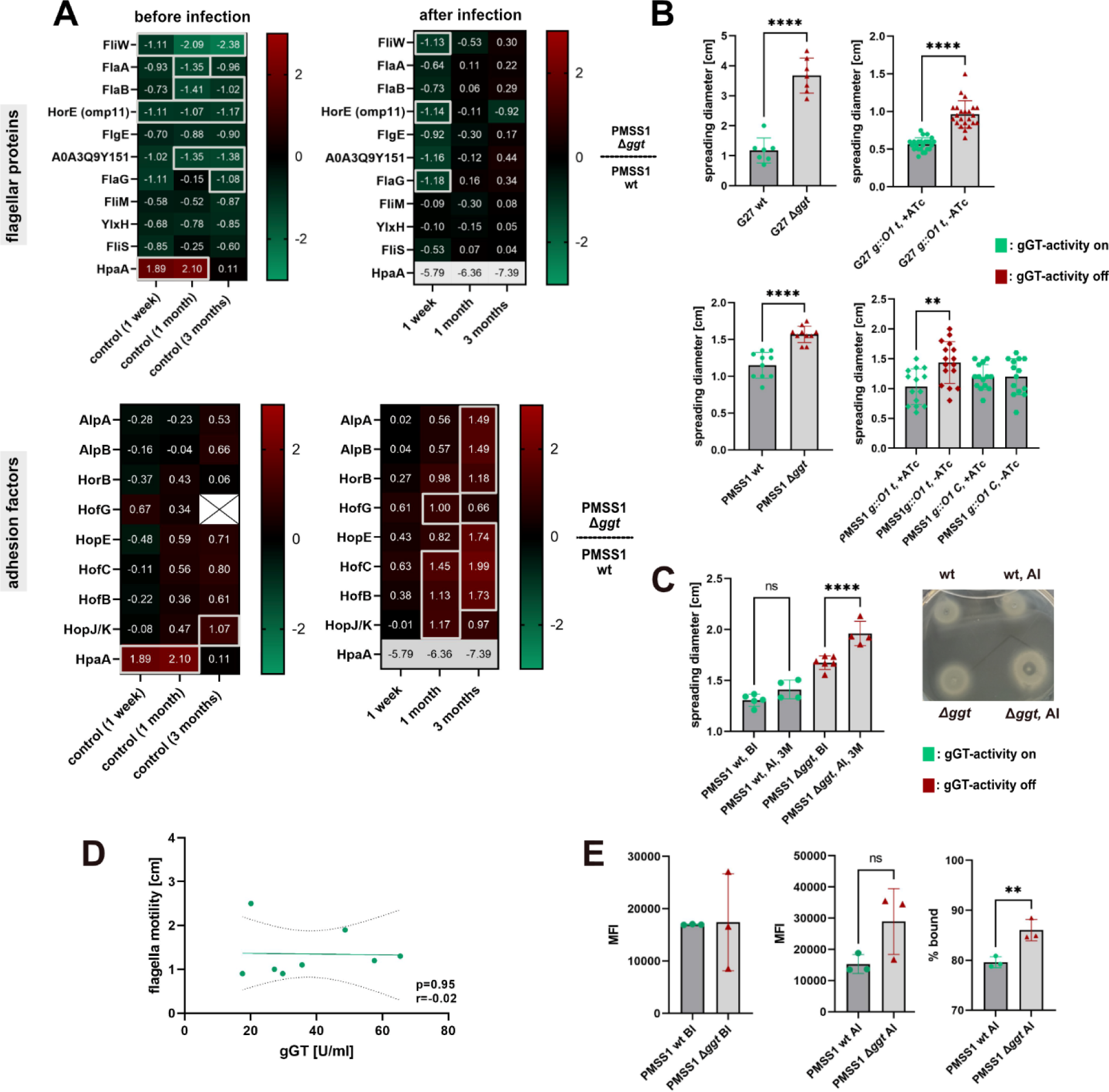
*H. pylori*’s motility and adhesion capacity is increased in the absence of gGT during infection. (**A**) Log_2_ transformed fold change of flagellar proteins and adhesion factors comparing the proteome of PMSS1 wt and Δ*ggt*. The proteomes of three single clones isolated after 1 week, 1 month and 3 months of infection from different mice were determined. Significant differences (-log_10_(p-value) > 1.3 and log_2_(fold change) ≥ I1I) are marked with grey boxes. Missing data points are colored white and are marked with a cross. Values out of the range of the heat map are colored grey. (**B and C**) Flagellar motility of *H. pylori* strains in the presence and absence of gGT. Soft agar assays on Brucella/10% FCS plates were performed. 100 ng/ml ATc was added to the plates to switch on gGT-activity in inducible strains. gGT-activity was not induced in the pre-culture prepared for the experiment. Motility of deletion strains and inducible strains was determined (B) as well as motility before (BI) and after infection (AI) (C). Bars represent the mean of 8 to 14 independent experiments as indicated by single dots (B) or the mean motility of 4 to 5 isolates as measured in three independent experiments (C). (**D**) Correlation of flagellar motility of clinical isolates with gGT-activity. Each dot represents the mean motility of an isolate as determined by 3 independent experiments (**E**) Adhesion capacity of *H. pylori* before infection (BI) and after infection (AI) to AGS cells. Bars represent the mean of three independent experiments. MFI is the median fluorescence intensity on the PE channel. Error bars represent the standard deviation. Student’s t-test (A, B, E). ANOVA (C), Person’s correlation (D). \**\** p<0.01, ****p<0.001.

After infection, substantial variations in flagellar proteins between wt and gGT-deficient strain were observed only in isolates obtained after one week of infection, but not in re-isolates from later infection time points. This suggests that expression or secretion of flagellar proteins shifts during the course of infection in gGT-deficient mutants that are able to colonize mice. These flagellar proteins included the flagellar assembly factor *FliW*, FlaA/B, Flagellar hook protein (FlgE), Flagellin G (FlaG) and omp11, which was hypothesized to be specific for the flagellar sheath membrane [41]. Notably, only the expression of the flagellar sheath associated protein HpaA displayed a distinct pattern, while expression of other flagellar proteins followed the same trend during infection. Increased levels of HpaA expression were detected prior to infection in the gGT-deficient mutant, which was found to be significantly decreased (60-fold) in gGT-deficient reisolates. We confirmed these results by Western blot (**S9 Fig**). Here, PMSS1 wt exhibited stronger expression of HpaA before infection, while its expression was reduced in re-isolates. In contrast, the wt strain showed lower expression of HpaA compared to isogenic gGT-deficient strains after infection.

To investigate whether the observed differences in flagellar proteins affected motility, soft agar assays were performed. Strikingly, deletion or silencing of gGT in both G27 and PMSS1 strains significantly increased motility (**Fig 5 B**). Specifically, motility of the gGT-deficient G27 strain was 3-fold higher compared to the isogenic gGT-proficient counterpart, while motility of PMSS1 increased 1.5-fold in the absence of gGT. In a next step, we assessed the effect of infection-induced changes in flagellar protein expression on motility. Therefore, motility of re-isolates was compared to the strains initially used for infection. The motility of PMSS1 wt remained unaltered after infection, whereas the gGT-deficient re-isolates gained motility in comparison to the strain initially used for infection (**Fig 5 C**). To determine whether gGT-activity correlates with motility, flagellar movement of clinical isolates with different gGT-activities was determined. Our findings suggest that there is no correlation between gGT-activity and flagellar motility and that other factors might play a more dominant role in controlling bacterial movement (**Fig 5 D**).

Collectively, these findings suggest that absence of gGT forces *H. pylori* to differentially regulate flagellar proteins, resulting in an increased motility of gGT-deficient strains, a trait that becomes even more pronounced during infection.

Furthermore, the absence of gGT during infection caused the emergence of re-isolates characterized by an increased expression of adhesion factors. While AlpA, AlpB [42] and HorB [43] are known to support *H. pylori* binding to gastric epithelial cells, other outer membrane proteins from the Hof- and Hop family including HofG, HopE, HofC, HofB and HopJ/K, affected by the absence of gGT, are only predicted to contribute to binding [44]. In comparison to the wt strain, only putative adhesins HpaA and HopJ/K displayed increased expression in the gGT-deficient mutant prior to infection, while re-isolates retrieved after 1 month or 3 months of infection exhibited substantially elevated expression levels of several known and putative adhesion factors (**Fig 5 A**). To dissect whether the mass-spectrometry detected variations result in alterations of the binding capacity of *H. pylori*, TAMRA-stained bacteria were added to AGS cells and the bacterial signal on AGS cells was quantified using flow cytometry. While strains initially used for infection displayed comparable binding profiles independently of gGT, gGT-deficient re-isolates exhibited an increased binding capacity to the gastric epithelial cell line as compared to wt re-isolates. In summary, these findings indicate that the absence of gGT during infection favors the selection of clones with higher adhesion capabilities (**Fig 5 E**)

Taken together, comparative analysis of the proteome of wt and gGT-deficient PMSS1 revealed that gGT-deficiency strongly prompts *H. pylori* to differentially express various bacterial proteins, encompassing metabolic enzymes, flagellar proteins, and adhesion factors.

## Discussion

*H. pylori* possesses a repertoire of virulence factors that enables its survival in the hostile stomach environment, allowing for persistent colonization of the gastric mucosa. In this study, we investigated the highly conserved virulence factor gGT and linked its activity to other bacterial traits important for successful colonization including acid survival, iron acquisition, motility, and adhesion.

Several studies have indicated the contributory role of *H. pylori* gGT to initial colonization [8, 11, 12]. However, the specific functional roles of the enzyme underlying its effect on initial colonization have remained elusive. Since an effect of gGT on the immune system was ruled out in this context [8], as colonization differences emerge prior to the infiltration of innate or adaptive immune cells, a metabolic advantage provided by the enzyme became a plausible explanation for the reduced colonization ability of strains deficient for gGT.

We found evidence that gGT contributes to bacterial survival in physiologic fasted state-like low pH conditions by supporting the function of the key enzyme urease. Although Miller and Maier [14] proposed a role of gGT for acid resistance, our study extends their findings by simulating different physiological acidic environments and, more importantly, by providing a new mechanism for the contribution of gGT to acid resistance. However, the functional relationship between gGT and urease activity is still unknown. Miller and Maier hypothesized that urea-derived ammonia was assimilated in the bacterial cytoplasm into nitrogen-containing molecules that are deaminated in the periplasm. In the absence of gGT, deamination of glutamine in the periplasm is abolished, potentially affecting ammonia assimilation in the cytoplasm, which would result in high concentrations of toxic ammonia and might ultimately cause *H. pylori* to limit its urease activity. Moreover, nickel is an essential cofactor for urease function [45]. Considering that host factors like lactoferrin can bind nickel in addition to iron [46] and that *fecA3* might be involved in the uptake of complexed nickel ions [47], an effect of gGT on nickel uptake also seems plausible which might affect urease function in gGT-deficient strains.

In addition to refining the role of gGT in acid resistance *in-vitro*, we were able to directly link gGT-dependent acid resistance to initial colonization *in-vivo*, confirming a pivotal role of gGT in the acidic stomach environment. In humans *H. pylori* seems to depend even more on gGT for infection than in the mouse model. While all human isolates express this enzyme [13] mouse infections with gGT deficient strains are possible [8, 12]. This intriguing difference may be attributed to the markedly lower pH levels in the human stomach, which are reported to be pH 1.7 in the fasted state [48], whereas the fasted murine stomach maintains a relatively less acidic pH of 4 [49].

Apart from being important to cope with acidity and thereby supporting initial colonization, it is very likely that additional functions and molecular mechanisms related to gGT activity play a role in colonization and persistence because, even after elevation of the stomach pH, the gGT-deficient bacteria were still not able to colonize at the same level than the wt and over time, reduced bacterial load was observed in mice infected with the gGT knockout strain. Those alternative mechanisms include alterations in important metabolic pathways, changes in flagellar proteins as well as differential regulation of molecules important for adherence to the gastric epithelium.

During infection, the host actively restricts iron availability for pathogens as part of the innate defense to limit growth of invading microbes [50]. *H. pylori* genes essential for iron acquisition, like *feoB* [40] or *fecA1* [51], for iron storage like *pfr* [52] or for the regulation of the iron metabolism, like *fur* [53], were reported to strongly support *H. pylori* colonization. Notably, we observed that gGT-deficient mutants are highly sensitive to iron-restricted environments, making it very likely that a hampered iron acquisition in the absence of gGT would be another mechanism contributing to the reduced colonization capacity of the gGT-deficient strain.

Although a direct functional link of gGT to iron assimilation has not been described for *H. pylori* yet, *H. pylori* gGT is predicted to be Fur regulated and thus involved in the iron metabolism of the bacterium [39]. Pich et al. already proposed a plausible hypothesis for the function of *H. pylori* gGT in iron acquisition based on findings in the mycopathogen *Histoplasma capsulatum* [38]. According to this hypothesis, Cys-Gly, produced by gGT-activity, reduces Fe^3+^, which is typically bound in complex molecules, resulting in soluble Fe^2+^ ions that can be taken up by *H. pylori*. This mechanism was already shown to enable *H. capsulatum* to use hemin and transferrin as iron sources because gGT-dependent production of Cys-Gly causes the release of iron from these sources [54, 55]. Nevertheless, further studies are needed to link the contribution of this mechanism to *H. pylori* colonization.

Apart from iron, the nitrogen metabolism is also crucial to support *H. pylori* growth in the stomach. *In-vitro* studies reported gGT to be important for the bacterial nitrogen metabolism by hydrolyzing glutamine which is followed by the uptake of glutamate via the transporter GltS [13]. The discovery of Kavermann et al. demonstrating the crucial role of this transporter for *H. pylori* colonization of Mongolian gerbils [56], strongly suggests that this gGT-dependent pathway is also important *in-vivo*. In addition, the dependency of *H. pylori* on this metabolic pathway *in-vivo* is supported by the observed upregulation of ammonium producing enzymes and peptide transporters that might help the bacterium to acquire nitrogen containing molecules in the absence of gGT by other means than glutamine hydrolysis.

Pathogens like *F. tularensis* [16] and *N. meningitidis* [17] were found to substantially profit from gGT-derived Cys-Gly in their intracellular growth. More recently, *H. pylori* was also shown to take up Cys-Gly in a gGT-dependent manner [15]. Our observation of a strong selection for the Cys-Gly specific transporter CptA in gGT-deficient mutants capable of colonizing the murine stomach, suggests that *H. pylori* could potentially profit from this gGT-dependent peptide during infection.

Additionally, nutrient limitation can trigger flagellar synthesis [57], while a low cellular energy status can activate chemotaxis via the sensor TlpD [58]. Consequently, potential nutrient or energy shortages in the absence of gGT might increase bacterial motility, enabling *H. pylori* to actively search for alternative nutrient sources and supporting thereby bacterial colonization. The fact that several proteins involved in flagellar *H. pylori* motility including the flagellar proteins flagellin A (FlaA), flagellin B (FlaB) [59] and FlgK [60] and the flagellar motor protein (MotB) [61] were already shown to be essential to attain robust colonization, further stresses the importance of this bacterial trait during infection.

In parallel, the increased levels of adhesion factors observed in the absence of gGT would favor a tighter bacterial adhesion to host cells. Although an upregulation of flagellar proteins and adhesins at the same time seems counterintuitive, one may hypothesize that the absence of gGT might force the bacterium to move to other infection sites to acquire nutrients, and as result *H. pylori* needs to interact with the epithelium more frequently. The presence of the adhesion factors AlpA, AlpB and HorB are well-known to be beneficial for *H. pylori* colonization [43, 62], suggesting that an increase in their expression during infection, as observed here, can be supportive for bacterial colonization. In addition, gGT-dependent alterations in the protein HpaA, as detected in this study, can also impact colonization levels because HpaA was found to be essential for *H. pylori* infection of mice [63] and Mongolian gerbils [56].

As pointed out above, *H. pylori* gGT also impacts bacterial persistence in the stomach. In bacteria like *F. tularensis*, gGT activity has been reported to enable the growth of the bacterium in macrophages by supplying the pathogen with free cysteine from glutathione, while the *in vitro* growth of the pathogen is not affected by the lack of the enzyme [16]. Considering this, it would be interesting to determine whether gGT activity represents an additional mechanism of *H. pylori* to survive in phagocytes. Moreover, gGT-activity induces the infiltration of CD8^+^ T cells, which were recently found to efficiently control *H. pylori* infection upon infiltration [64]. Thus, it will be of future interest to analyze whether these immune cells might be involved in gGT-dependent *H. pylori* persistence *in-vivo*.

This study uncovered several novel mechanistic functions of *H. pylori* gGT through the analysis of re-isolates from a mouse infection model. Although findings were validated with clinical isolates, indicating their relevance in the human context, it is important to note that using a mouse model may potentially limit the functional mechanisms of gGT that can be detected. Conditions in the human stomach vary from those in the murine stomach, including differences in terms of acidity [48, 49] anatomy [65] and the availability of nutrients from food intake. This suggests that *H. pylori* might face distinctive selective forces in the human host compared to the mouse model resulting in differently adapted isolates. Moreover, by analyzing re-isolates only long-term adaptations can be detected, potentially overlooking short-term changes in protein expression that might occur in a complex and changeable environment such as the stomach.

In summary, *H. pylori* gGT has a pivotal role in acid resistance and iron acquisition, while its loss induces several compensatory mechanisms to ensure bacterial colonization and persistence in the host. These findings may have potential implications for drug development. On the one hand, *H. pylori* gGT is a promising target for the development of new therapies because the enzyme is functionally highly important for the bacterium on different levels such as immune evasion, acid resistance and nutritional acquisition. On the other hand, our results suggest, that targeting *H. pylori* gGT alone may not be sufficient to limit bacterial colonization, as *H. pylori* can compensate for the loss of the enzyme. Therefore, the combination with other targets identified in our study as potential compensatory candidates might mitigate the emergence of escape mutants.

## Materials and Methods

### *H. pylori* strains and culture conditions

*H. pylori* was grown in microaerobic conditions at 37 °C, 10% (*v*/*v*) CO_2_ and 5% (*v*/*v*) O_2_ on WC sheep blood agar plates supplemented with Dent (Oxoid) or in BHI medium supplemented with 10% FCS shaking at 120 rpm. In this study, the *H. pylori* laboratory strains G27 [66], and PMSS1 [67] as well as clinical isolates were used. PMSS1 Δ*ureAB* was generated by transformation of genomic DNA from G27 Δ*ureAB* [68]. Mutant strains were grown with antibiotic selection in the growth medium (50 µg/ml kanamycin, 10 µg/ml streptomycin or 10 µg/ml chloramphenicol). All strains used in this study are listed in S1 Table. To induce gGT-activity in *H. pylori g::O1 t*, 100 ng/ml anhydrotetracycline (ATc) was added to the culture for 24 h, if not stated otherwise. If *H. pylori g::O1 t* was used for experiments, gGT-activity was induced in the pre-culture and during the experiment except as otherwise indicated. In some experiments *H. pylori* was grown in acidic Brucella medium (pH 5) supplemented with 20% FCS. The pH was adjusted with hydrogen chloride and pH was monitored during the experiment with a portable pH measuring device (FiveGo pH meter F2-Std-Kit, Mettler-Toledo).

### Construction of conditional *H. pylori* gGT mutants

Conditional gGT mutants were constructed using a counter selection system based on a *rpsL-cat* cassette as described by Dailidiene et al. and Debowski et al. [20, 69]. First, streptomycin resistant *H. pylori* strains were generated by replacing *H. pylori rpsL* by a mutated version. Plasmids containing either *rpsL-cat* or the inducible promoter *tetO1* [18] flanked by regions homologous to the DNA up- and downstream of the *ggt*-promoter were constructed using a commercially available Gibson Assembly Cloning Kit (NEB). Homologous regions were amplified from the genomic DNA of strain PMSS1 using primer pairs FlgK fwd/FlgK rev (rpsL/tetO1) and ggt fwd (rpsL/tetO1)/ggt rev. The *rpsL-cat* cassette and the *tetO1* promoter were amplified from pOND708 and pOND817 [18] by the primers rpsL fwd/rev and tetO1 fwd/rev, respectively. pOND708 was opened by PCR using the primers backbone (rpsL/tetO1) fwd/rev and was used as a plasmid backbone. An overview of the cloning procedure (S1 Fig), primers (S2 Table) and plasmids (S3 Table) is presented in the supplemental material. Plasmids were naturally transformed to streptomycin resistant *H. pylori* to replace the *ggt* promoter by *tetO1*. Transformants were tested for correct genomic insertions by PCR using the primers FlgK fwd seq and tetO1 rev. Next, natural transformation of pOND708 and pOND1162 [18] resulted in the introduction of *tetR* into the *H. pylori* genome. Correct insertion was verified by PCR using the primer pair TetR check fwd/rev.

### Mouse experiments

Female C57BL/6 wild-type mice (Envigo) were housed under specific pathogen-free conditions and were fed *ad libitum*. Six- to eight-week-old mice were infected with *H. pylori* PMSS1 wt or with PMSS1 Δ*ggt* by oral gavage with 2*10^8^ bacteria resuspended in 200 µl BHI with 20% FCS. Mice were challenged twice with *H. pylori* at a time interval of two days. A group of mice received 200 µl sodium bicarbonate (10% w/v) 10 min before infection to elevate the pH in the murine stomach. Colonization was analyzed by plating serial dilutions of a longitudinal piece of stomach homogenized in BHI/20% FCS on WC-dent blood agar plates supplemented with bacitracin (200 μg/ml), nalidixic acid (10 μg/ml), and polymyxin B (3 μg/ml). After 5 days of culture, single clones were counted and expanded on WC-dent plates for 2 days and then either frozen for later use or directly grown in liquid culture for proteomic analysis. All experiments were approved by the Bavarian Government (Regierung von Oberbayern, ROB-55.2-2532.Vet_02-19-95) and conducted in compliance with European guidelines for the care and use of laboratory animals.

### Proteomics

*H. pylori* strains were grown to the early stationary growth phase or for a maximum of 35 h, washed twice with PBS, and lysed in RIPA buffer (50 mM Tris, 150 mM NaCl, 1 mM EGTA, 1% Igepal, 0.25% sodium deoxycholate, pH 7.4) using ultrasonication. The insoluble fraction was separated by centrifugation (30 min, 10 000 g, 4 °C) and protein concentration in the supernatant was determined by a commercially available BCA assay (ThermoFisher Scientific). The same protein amount was used for each measurement.

Proteins were precipitated with ice-cold acetone (−20 °C, overnight), centrifuged (21 000 g, 15 min, 4 °C) and dissolved in cold methanol (−80 °C) using sonication (10 s, 10%, 5x cycle). Proteins were resuspended in denaturation buffer (7 M urea, 2 M thiourea in 20 mM HEPES, pH 7.5) and reduced in the presence of 10 mM TCEP (1 h, 37 °C, 600 rpm). Alkylation was performed with 10 mM IAA (30 min, 1 h, 600 rpm) and was quenched by adding DTT to a final concentration of 10 mM (30 min, 600 rpm). Proteins were pre-digested with LysC (Promega) for 2 h at 25 °C and digested with trypsin (Promega, 37 °C, overnight) in TEAB buffer (50 mM, triethylammonium bicarbonate). Digestion was stopped by acidification with formic acid to a pH below 3. SepPak C18 cartridges (Waters) were used for desalting. Cartridges were equilibrated with 1 ml elution buffer (80% acetonitrile, 0.5% formic acid) and 0.1% TFA, before peptides were loaded on the column. Washing was done with 0.1% TFA (2x 1 ml) and with 0.5% formic acid (1x 0.5 ml). Peptides were eluted with elution buffer (1x 500 μl, 1x 250 μl) and dried by lyophilization. After reconstitution in 1% formic acid to a final concentration of 1 µg/µl, peptides were filtered through centrifugal filter units (Merck, PVDF, 0.22 µm) before being transferred to MS vials.

MS analysis was performed on an Orbitrap Fusion mass spectrometer coupled to Ultimate3000 nano-HPLC via a *Nanospray Flex* Ion Source (ThermoFisher Scientific). Samples were loaded on the trap column (AcclaimPepMap 100 C18 (75 μm×2 cm) trap/ flow rate: 5 µl/min, 0.1 % TFA) and separated on the Aurora Ultimate^TM^ columns (2^nd^ generation, 75 µm×25 cm, *ionopticks*). Both columns were constantly heated to 40 °C. For separation, a buffer B gradient (0.1% formic acid in acetonitrile) with a flowrate of 400 nL/min was applied (5-22% buffer B for 112 min, to 32% buffer B in 10 min, to 90% buffer B in 10 min, hold for 10 min, to 5% buffer B in 0.1 min, hold 5% buffer B for 9.9 min). The Orbitrap was operated in a cycle time (3 s) data dependent mode. An AGC target of 2e5, a maximum injection time of 50 ms, 60% RF lens and a resolution of 120 000 in a scan range of 300-1500 *m/z* in profile mode was used. Monoisotopic precursor selection and dynamic exclusion (60 s) was turned on. For fragmentation, most intense precursors with charges of 2-7 and intensities greater than 5e3 were chosen. Quadrupole isolation was performed using a range of 1.6 *m/z.* Precursor ions were separated using an AGC target of 1e4 and a maximum injection time of 35 ms. Fragmentation was performed using higher-energy collisional dissociation with a collision energy of 30%. Fragments were detected in the ion trap operating at a rapid scan rate.

Data acquisition was done using Xcalibur software followed by data processing using MaxQuant 1.6.2.10. [70]. Proteins were searched against the data available for *H. pylori* PMSS1 (UP000289502) or G27 (UP000001735) in the UniProt database. Further bioinformatical analysis was performed with Perseus version 1.6.14.10 [71]. Potential contaminants and reverse hits were removed. Label-free quantification intensities were log_2_-transformed and proteins with less than three valid values were filtered off. Missing values were replaced by imputed numbers from a matrix (width 0.3, downshift 1.8, total matrix). To identify relevant differences Student’s t-test was used. Hits with a *p*-value < 0.05 and a Student’s t-test difference of at least |1| were considered to be significant. Proteomics data can be accessed via the PRIDE repository with the identifier PXD050334.

### Urease, amidase and formamidase activity assays

Enzymatic activities of urease [72], AmiE and AmiF [73] were determined as previously described with some modifications.

*H. pylori* (4*10^8^ bacteria/ml) was harvested in PEB (100 mM sodium phosphate buffer, pH 7.3, 10 mM EDTA), washed once with the same buffer and lysed by 10 min sonication. Lysates were cleared from cell debris by centrifugation (10 000 *g*, 10 min). To measure urease activity after exposure to an acidic environment, bacteria were resuspended in McIlvaine buffer (pH 5 or pH 7) first and incubated for 2 h prior to lysis. For measurement of enzymatic activity in the supernatant, supernatant after 2 h of incubation with bacteria was collected. Protein concentration of lysates and supernatants was determined by a commercially available BCA assay (ThermoScientific).

Samples (90 µl) were mixed with 100 µl of reaction buffer (50 mM urea or 100 mM acrylamide or 1 mM formamide in PEB). The hydrolysis of urea was stopped after 10 min at room temperature, the hydrolysis of formamide and acrylamide after 30 min at 30 °C by the addition of 100 µl phenol-nitroprusside (Sigma). For color development 50 µl alkaline hypochlorite (Sigma) were added and absorption was measured after 1 h (Tecan plate reader, 670 nm). The amount of ammonia released was determined from a standard curve with ammonium chloride (0-500 µM). One unit (U) of enzymatic activity was defined as the amount of enzyme that hydrolyses one µmol of substrate per minute under the conditions of the assay.

### gGT activity assay

gGT-activity was determined by a colorimetric assay based on the release of p-nitroaniline from the substrate gGpNA (Sigma). The method was adapted from Meister et al. (1981) [74] and from Schmees et al. [75] with some modifications. *H. pylori* was resuspended in PBS (2* 10^8^ bacteria/ml) and incubated for 2 h in microaerobic conditions shaking at 120 rpm. Supernatant and bacteria were separated by centrifugation (5000 *g*, 5 min) and 50 µl of the supernatant was used for the assay. Supernatant and 150 µl reaction buffer (5 mM gGpNA, 100 mM Gly-Gly in 0.1 M Tris buffer, pH 8) were mixed. After 1 h of incubation at 37 °C the release of p-nitroaniline was measured (Tecan plate reader, 405 nm). One unit (U) of activity was defined as the amount of enzyme that releases one µmol of p-nitroanilide per minute under the conditions of the assay.

### Growth in iron-restricted medium

0.2*10^8^ bacteria/ml were resuspended in BHI supplemented with 10% FCS and with deferoxamine mesylate (DFO) at different concentrations (0 µM, 20 µM, 50 µM or 75 µM). Bacteria were grown for 48 h. Growth was measured spectrophotometrically at 600 nm using a BioPhotometer (Eppendorf). Spectrometric growth measurements were normalized to bacterial growth without DFO.

### Acid survival assay

Acid survival was measured by the method described by Miller and Maier with some modifications [14]. *H. pylori* was grown in liquid culture for 24 h, harvested and washed with PBS once. Bacteria were resuspended in 20 ml of acid challenge buffer at a concentration of 0.2*10^8^ bacteria/ml. McIlvaine buffer (pH 3 or pH 5) or fasted state simulated gastric fluid (FaSSGF, [27]) were used. For some experiments acid challenge buffers were supplemented with 8 mM urea. Bacteria were incubated in microaerobic conditions at 37 °C shaking at 120 rpm. Survival was measured by plating serial dilutions on WC-dent blood agar plates.

### Soft agar motility assay

*H. pylori* motility was assessed as previously described [76]. The soft agar plates consisted of Brucella broth, 0.35% (w/v) agar, 10% (v/v) FCS and *H. pylori* selective supplement (Dent). Using a sterile toothpick, *H. pylori* was inoculated into the soft agar. Migration of the bacteria was monitored for up to five days by measuring the diameter of the bacterial halo around the starting point.

### Flow cytometry based binding assay

AGS cells (ATCC CRL-1739) were cultured in DMEM (Gibco) containing 10% FCS and 1% Pen-Strep (Gibco, 5000 U/ml) at 5% CO_2_ at 37 °C. A *H. pylori* culture was adjusted to a concentration of 2*10^8^ bacteria/ml in BHI supplemented with 20% FCS and was stained with TAMRA (Sigma, 1 µl TAMRA/ml) for 30 min shaking at 120 rpm in microaerobic conditions at 37 °C. AGS cells were detached with trypsin (Gibco), resuspended in antibiotic-free DMEM and stained with DAPI (Sigma, 3 µl DAPI/ml) for 30 min at 37°C. Bacteria and cells were washed twice with PBS to remove excessive dye. AGS cells were seeded in a 96-Well plate (1*10^5^ cells/well) in antibiotic-free DMEM supplemented with 10% BHI/20%FCS, mixed with bacteria at a MOI of 5 and medium was topped up to 200 µl. After 30 min at 37 °C, unbound bacteria were washed off with PBS in three washing steps. Cells were fixed with 1% PFA in PBS, washed with PBS and resuspended in FACS buffer (0.5% BSA in PBS). Binding was analyzed by flow cytometry (Cytoflex S, Beckman Coulter) by comparing the TAMRA-signals (PE-channel) of AGS cells.

### Statistical analysis

Results of *in-vitro* experiments are presented as mean ± standard deviation (SD) of three independent experiments, if not stated otherwise. Results from *in-vivo* experiments are presented as dot plots with medians. The normal distribution of data was tested using the Shapiro-Wilk test. Normally distributed data was analyzed using a t-test or one-way ANOVA followed by multiple Bonferroni corrected comparisons. For non-normally distributed data, Mann-Whitney U-Test was used. Statistical significance was established when p< 0.05, while highly statistically significant results had a p-value of p< 0.01.

## Acknowledgements

We thank the biopharmaceutical company Ondek for providing the plasmids pOND708, pOND817 and pOND1162 that were used to create the tet system presented in this work. The cartoon in Fig 1 A and in S1 Fig was created with BioRender.com.

## Supporting information captions

**S1 Table: Strains used in this study**

**S1 Table: Oligonucleotide primers used in this study**

**S3 Table: Plasmids used in this study**

**S4 Table: Significantly different proteins comparing the proteome of PMSS1 wt re-isolates with PMSS1 Δ*ggt* re-isolates**

**S2 Table: Significantly different proteins comparing the proteome of PMSS1 wt with PMSS1 Δ*ggt***

**S3 Table: Significantly different proteins comparing the proteome of PMSS1 Δ*ggt* re-isolates with the proteome of the PMSS1 Δ*ggt* isolate used for infection**

**S7 Table: Significantly different proteins comparing the proteome of PMSS1 wt re-isolates with the proteome of the PMSS1 wt isolate used for infection**

**S1 Fig. Cloning procedure**

Plasmid backbone and rpsL-cat were amplified by PCR from the plasmid pOND708 using the primer pairs backbone fwd/rev and rpsL fwd/rev, respectively. Homologous flanks called FlgK flank and ggt flank were retrieved from the genomic DNA of PMSS1 by PCR with the primers Flgk flank fwd/rev and ggt flank fwd/rev. A plasmid (pGGT_rpsL-cat) to exchange the *ggt* promoter with *rpsL-cat*, was assembled using the plasmid backbone, homologous flanks and *rpsL-cat*. The plasmid pGGT_tetO1 was generated similarly but instead of the rpsL-cat cassette the *tetO1* promoter was used for Gibson Assembly. This promoter was amplified before the assembly from pOND817 using the primers tetO1 fwd and rev.

**S1 Fig. PCR confirmation of mutant strains used in this study**

PCRs using GoTaq Green Master Mix (Promega) were performed to confirm the genotype of the strains used in this study. Genomic DNA was purified before PCR using a commercially available kit (GenUP™ Bacteria gDNA Kit, biotech rabbit). Primer pairs ggt fwd/rev, ureAB fwd/rev, TetR seq fwd/rev and FlgK seq fwd/tetO1 rev (**C**) were used to detect the presence of *ggt* (**A**)*, ureAB* (**B**), *tetR* and *tetO1* (**C**) in the *H. pylori* genome, respectively. Benchtop 100 bp or 1 kb ladder (Promega) was used.

**S3 Fig. Establishment and characterization of inducible tet-on gGT *H. pylori* strains**

(**A**) Western Blot detecting *tetR*-controlled induction of gGT-expression in PMSS1 *g::O1 t.* For induction, 100 ng/ml ATc was added to the culture. As a control lysate of PMSS1 *g::O1 t* after induction for 24 h was used. Equal amounts of protein (2 µg) were loaded in each lane. (**B**) Growth of *H. pylori* wt in the presence of different ATc concentrations. Growth was monitored spectrophotometrically at 600 nm. ATc was dissolved in ethanol. As solvent control the same amount of ethanol was added to each culture. Shown is one representative experiment per strain. n=3 (**C**) TetR controlled induction and silencing of gGT-activity in G27 *g::O1 t*. For induction, 100 ng/ml ATc was added to the culture (starting concentration: 1×10^8^ bacteria/ml). As a control the gGT-activity of G27 *g::O1 t* after induction for 24 h was measured. For silencing, gGT-activity of G27 *g::O1 t* was induced for 24 h with 100 ng/ml ATc during growth in BHI/10% FCS. The culture was washed with PBS and resuspended in ATc-free medium at a concentration of 1*10^8^ bacteria/ml. As a control, an uninduced 24 h old culture of G27 *g::O1 t* was used. Bars represent the mean of one or two independent experiments as represented by single dots. n: number of independent experiments.

**S4 Fig. Growth and pH increase of *H. pylori* at low pH**

(**A**) Growth of *H. pylori* in BHI medium with 20% FCS at pH 5 and pH 7. Growth was measured spectrophotometrically at 600 nm (OD_600_). One representative experiment is shown. (n=2) (**B**) pH increase during growth in Brucella medium supplemented with 20% FCS at pH 5 and pH 7. PH was measured with a handheld electrode. (n=3). n: number of independent experiments.

**S2 Fig. *H. pylori* gGT contributes to acid survival**

(**A, B and C**) Survival of *H. pylori* in McIlvaine buffer without supplementation at pH 5 (A) or pH 3 (B) and in fasted state simulated gastric fluid (FaSSGF) (C), viable cell counts were monitored by plating. Shown are the means and range of three independent experiments (A and B) or one representative experiment (n=3) (C). The detection limit of the assay is indicated by a dotted line. (**D**) *H. pylori* induced urea-dependent pH increase. Bacteria were grown on WC-dent plates and resuspended in McIlvaine buffer (4*10^8^ bacteria/ml) pH 5 with 8 mM urea. PH was monitored with a handheld pH electrode. (**E**) Urease activity in the supernatant after exposure of *H. pylori* for 2 h to pH 5. Strains were grown in BHI with 10% FCS for 24 h and resuspended for 2 h in McIlvaine buffer (pH 5) before measurement. Data was normalized to the mean activity of *H. pylori* wt. Bars represent the mean of 4-5 independent experiments as indicated by single dots. Error bars represent the standard deviation. (**F**) Western blot detection of urease subunit B in the supernatant and lysate. The same amounts of protein were loaded (supernatant: 0.5 µg, lysate: 5 µg). CagA was used as a loading control. n: number of independent experiments. ANOVA (D, E).

**S6 Fig. Comparison of the proteome and secretome of *H. pylori* wt and Δ*ggt* after exposure to pH 5.**

The proteomes and secretomes of *H. pylori* wt and Δ*ggt* were determined in triplicates. Thresholds for significance (-log_10_(p-value) > 1.3 and log_2_(fold change) ≥ I1I) are represented by dotted lines. Proteins stronger expressed by the wt are marked in green, proteins stronger expressed by the gGT-deficient mutants are marked in red. Flagellar proteins are highlighted in orange. Students t-test.

**S3 Fig. Preparation for proteome comparison of *H. pylori* isolates**

**(A)** Infection level of mice used for re-isolation of *H. pylori*. C57Bl6 mice were inoculated twice with 2*10^8^ bacteria at a time interval of 2 days. Mice were sacrificed after 1 week, 1 month of 3 months of infection and colonization level was assessed by plating stomach homogenates. **(B)** Growth curves of re-isolates and strains initially used for infection (BI). Isolates were picked, expanded for 2 d on WC-dent plates and cultivated in BHI/10% FCS for a maximum of 35 h or until the stationary growth phase was reached. Mann Whitney U-test. *p < 0.05, **p < 0.01

**S8 Fig. Formamidase and iron uptake activity is increased in the absence of gGT**

(**A and B**) AmiE and AmiF activity of *H. pylori* strains before infection (BI) and of re-isolates (RI). Strains were grown for 24 h in BHI/10% FCS before measurement. RI were retrieved from mice stomachs after 1 month of infection, expanded for 2 d on plates and frozen for further use. (**C**) Fe(III) citrate uptake of wt and *ggt* deletion strains. Bars represent the mean of three to four independent experiments as indicated by dots. Error bars represent the standard deviation. ANOVA. *\** p<0.05.

**S9 Fig. Immunoblot detection of HpaA**

Bacteria were grown for two days on WC DENT plates and were lysed with RIPA buffer. 5 µg lysate and 0.5 µg recombinant HpaA were loaded on the SDS gel. CagA was used as a loading control and recombinant HpaA as a positive control.

## Additional information

### Financial disclosure statement

This work was supported by the FG under the grant GE 2042/11-1

### Data availability statement

All data generated or analyzed during this study are included in this article. Mass spectrometry proteomics data have been deposited to the ProteomeXchange [77] Consortium via the PRIDE [78] partner repository with the dataset identifier PXD050334. Other datasets are available from the corresponding author on reasonable request.

### PRIDE reviewer account

**Password:** tNyIllCa

### Competing interests

The authors report there are no competing interests to declare.

## Notes

### Competing Interest Statement

The authors have declared no competing interest.

## References

1. Malfertheiner P, Camargo MC, El-Omar E, Liou JM, Peek R, Schulz C, et al. Helicobacter pylori infection. Nat Rev Dis Primers. 2023;9(1):19.10.1038/s41572-023-00431-8

2. Rimbara E, Mori S, Kim H, Shibayama K. Role of gamma-glutamyltranspeptidase in the pathogenesis of Helicobacter pylori infection. Microbiol Immunol. 2013;57(10):665–73.10.1111/1348-0421.12089

3. Gong M, Ling SS, Lui SY, Yeoh KG, Ho B. Helicobacter pylori gamma-glutamyl transpeptidase is a pathogenic factor in the development of peptic ulcer disease. Gastroenterology. 2010;139(2):564–73.10.1053/j.gastro.2010.03.050

4. Shibayama K, Wachino J, Arakawa Y, Saidijam M, Rutherford NG, Henderson PJ. Metabolism of glutamine and glutathione via gamma-glutamyltranspeptidase and glutamate transport in Helicobacter pylori: possible significance in the pathophysiology of the organism. Mol Microbiol. 2007;64(2):396–406.10.1111/j.1365-2958.2007.05661.x

5. Megraud F, Neman-Simha V, Brugmann D. Further Evidence of the Toxic Effect of Ammonia Produced by Helicobacter pylori Urease on Human Epithelial Cells. INFECTION AND IMMUNITY. 1992;60(5):1858–63.10.1128/iai.60.5.1858-1863.1992

6. Matsui T, Matsukawa Y, Sakai T, Nakamura K, Aoike A, Kawai K. Effect of ammonia on cell-cycle progression of human gastric cancer cells. Eur J Gastroenterol Hepatol. 1995;7 Suppl 1:S79–81.10.1023/a:1018837920769

7. Oertli M, Noben M, Engler DB, Semper RP, Reuter S, Maxeiner J, et al. Helicobacter pylori gamma-glutamyl transpeptidase and vacuolating cytotoxin promote gastric persistence and immune tolerance. Proc Natl Acad Sci U S A. 2013;110(8):3047–52.10.1073/pnas.1211248110

8. Wustner S, Anderl F, Wanisch A, Sachs C, Steiger K, Nerlich A, et al. Helicobacter pylori gamma-glutamyl transferase contributes to colonization and differential recruitment of T cells during persistence. Sci Rep. 2017;7(1):13636.10.1038/s41598-017-14028-1

9. Wustner S, Mejias-Luque R, Koch MF, Rath E, Vieth M, Sieber SA, et al. Helicobacter pylori gamma-glutamyltranspeptidase impairs T-lymphocyte function by compromising metabolic adaption through inhibition of cMyc and IRF4 expression. Cell Microbiol. 2015;17(1):51–61.10.1111/cmi.12335

10. Kaebisch R, Semper RP, Wustner S, Gerhard M, Mejias-Luque R. Helicobacter pylori gamma-Glutamyltranspeptidase Induces Tolerogenic Human Dendritic Cells by Activation of Glutamate Receptors. J Immunol. 2016;196(10):4246–52.10.4049/jimmunol.1501062

11. Chevalier C, Thiberge J, Ferrero R, Labigne A. Essential role of Helicobacter pylori gamma-glutamyltranspeptidase for the colonization of the gastric mucosa of mice. Mol Microbiol. 1999;31(5):1359–72.10.1046/j.1365-2958.1999.01271.x

12. McGovern KJ, Blanchard TG, Gutierrez JA, Czinn SJ, Krakowka S, Youngman P. gamma-Glutamyltransferase is a Helicobacter pylori virulence factor but is not essential for colonization. Infect Immun. 2001;69(6):4168–73.10.1128/IAI.69.6.4168-4173.2001

13. Leduc D, Gallaud J, Stingl K, de Reuse H. Coupled amino acid deamidase-transport systems essential for Helicobacter pylori colonization. Infect Immun. 2010;78(6):2782–92.10.1128/IAI.00149-10

14. Miller EF, Maier RJ. Ammonium metabolism enzymes aid Helicobacter pylori acid resistance. J Bacteriol. 2014;196(17):3074–81.10.1128/JB.01423-13

15. Baskerville MJ, Kovalyova Y, Mejias-Luque R, Gerhard M, Hatzios SK. Isotope tracing reveals bacterial catabolism of host-derived glutathione during Helicobacter pylori infection. PLoS Pathog. 2023;19(7):e1011526.10.1371/journal.ppat.1011526

16. Alkhuder K, Meibom KL, Dubail I, Dupuis M, Charbit A. Glutathione provides a source of cysteine essential for intracellular multiplication of Francisella tularensis. PLoS Pathog. 2009;5(1):e1000284.10.1371/journal.ppat.1000284

17. Takahashi H, Hirose K, Watanabe H. Necessity of meningococcal gamma-glutamyl aminopeptidase for Neisseria meningitidis growth in rat cerebrospinal fluid (CSF) and CSF-like medium. J Bacteriol. 2004;186(1):244–7.10.1128/JB.186.1.244-247.2004

18. Debowski AW, Verbrugghe P, Sehnal M, Marshall BJ, Benghezal M. Development of a tetracycline-inducible gene expression system for the study of Helicobacter pylori pathogenesis. Appl Environ Microbiol. 2013;79(23):7351–9.10.1128/AEM.02701-13

19. Debowski AW, Sehnal M, Liao T, Stubbs KA, Marshall BJ, Benghezal M. Expansion of the tetracycline-dependent regulation toolbox for Helicobacter pylori. Appl Environ Microbiol. 2015;81(23):7969–80.10.1128/AEM.02191-15

20. Debowski AW, Walton SM, Chua EG, Tay AC, Liao T, Lamichhane B, et al. Helicobacter pylori gene silencing in vivo demonstrates urease is essential for chronic infection. PLoS Pathog. 2017;13(6):e1006464.10.1371/journal.ppat.1006464

21. Kalantzi L, Goumas K, Kalioras V, Abrahamsson B, Dressman JB, Reppas C. Characterization of the human upper gastrointestinal contents under conditions simulating bioavailability/bioequivalence studies. Pharm Res. 2006;23(1):165–76.10.1007/s11095-005-8476-1

22. Zhu H, Hart CA, Sales D, Roberts NB. Bacterial killing in gastric juice--effect of pH and pepsin on Escherichia coli and Helicobacter pylori. J Med Microbiol. 2006;55(Pt 9):1265–70.10.1099/jmm.0.46611-0

23. Han S, Evans D, el-Zaatari F, Go M, Graham D. The interaction of pH, bile, and Helicobacter pylori may explain duodenal ulcer. Am J Gastroenterol. 1996;91(6):1135–7

24. Schreiber S, Konradt M, Groll C, Suerbaum S. The spatial orientation of Helicobacter pylori in the gastric mucus. PNAS. 2004;101(14):5024–9.10.1073/pnas.030838610

25. Kim H, Park C, Jang WI, Lee KH, Kwon SO, Robey-Cafferty SS, et al. The gastric juice urea and ammonia levels in patients with Campylobacter pylori. Am J Clin Pathol. 1990;94(2):187–91.10.1093/ajcp/94.2.187

26. Marshall B, Barrett L, Prakash C, McCallum R, Guerrant R. Urea protects Helicobacter (Campylobacter) pylori from the bactericidal effect of acid. Gastroenterology. 1990;99(3):697–702.10.1016/0016-5085(90)90957-3

27. Jantratid E, Janssen N, Reppas C, Dressman JB. Dissolution media simulating conditions in the proximal human gastrointestinal tract: an update. Pharm Res. 2008;25(7):1663–76.10.1007/s11095-008-9569-4

28. Vertzoni M, Dressman J, Butler J, Hempenstall J, Reppas C. Simulation of fasting gastric conditions and its importance for the in vivo dissolution of lipophilic compounds. Eur J Pharm Biopharm. 2005;60(3):413–7.10.1016/j.ejpb.2005.03.002

29. Bucker R, Azevedo-Vethacke M, Groll C, Garten D, Josenhans C, Suerbaum S, Schreiber S. Helicobacter pylori colonization critically depends on postprandial gastric conditions. Sci Rep. 2012;2:994.10.1038/srep00994

30. Jiang X, Doyle MP. Effect of environmental and substrate factors on survival and growth of Helicobacter pylori. J Food Prot. 1998;61(8):929–33.10.4315/0362-028x-61.8.929

31. Bijlsma J, Lie-A-Ling M, Nootenboom I, Vandenbroucke-Grauls C, Kusters J. Identification of Loci Essential for the Growth of Helicobacter pylori under Acidic Conditions. The Journal of Infectious Diseases. 2000;182:1566–9.10.1086/315855

32. Scott D, Weeks D, Hong C, Postius S, Melchers K, Sachs G. The Role of Internal Urease in Acid Resistance of Helicobacter pylori. Gastroenterology 1998;114:58–70.10.1016/s0016-5085(98)70633-x.

33. Sachs G, Scott D, Weeks D, Melchers K. The importance of the surface urease of Helicobacter pylori: fact or fiction? Trends in Microbiology. 2001;9(11): 532–4.10.1016/S0966-842X(01)02226-0.

34. Phadnis S, Parlow M, Levy M. Surface localization of Helicobacter pylori urease and a heat shock protein homolog requires bacterial autolysis. Infect Immun 1996;64(3):905–12.10.1128/iai.64.3.905-912.1996

35. De Reuse H, Skouloubris S. Nitrogen Metabolism. In: Mendz GL HS, editor. Helicobacter pylori: Physiology and Genetics. Washington (DC): ASM Press; 2001.

36. Vorwerk H, Mohr J, Huber C, Wensel O, Schmidt-Hohagen K, Gripp E, et al. Utilization of host-derived cysteine-containing peptides overcomes the restricted sulphur metabolism of Campylobacter jejuni. Mol Microbiol. 2014;93(6):1224–45.10.1111/mmi.12732

37. Weinberg MV, Maier RJ. Peptide transport in Helicobacter pylori: roles of dpp and opp systems and evidence for additional peptide transporters. J Bacteriol. 2007;189(9):3392–402.10.1128/JB.01636-06

38. Pich OQ, Merrell DS. The ferric uptake regulator of Helicobacter pylori: a critical player in the battle for iron and colonization of the stomach. Future Microbiol. 2013;8(6):725-

38. 10.2217/fmb.13.43

39. Pich OQ, Carpenter BM, Gilbreath JJ, Merrell DS. Detailed analysis of Helicobacter pylori Fur-regulated promoters reveals a Fur box core sequence and novel Fur-regulated genes. Mol Microbiol. 2012;84(5):921–41.10.1111/j.1365-2958.2012.08066.x

40. Velayudhan J, Hughes NJ, McColm AA, Bagshaw J, Clayton CL, Andrews SC, Kelly DJ. Iron acquisition and virulence in Helicobacter pylori: a major role for FeoB, a high-affinity ferrous iron transporter. Mol Microbiol. 2000;37(2):274–86.10.1046/j.1365-2958.2000.01987.x

41. Josenhans C, Niehus E, Amersbach S, Horster A, Betz C, Drescher B, et al. Functional characterization of the antagonistic flagellar late regulators FliA and FlgM of Helicobacter pylori and their effects on the H. pylori transcriptome. Mol Microbiol. 2002;43(2):307–22.10.1046/j.1365-2958.2002.02765.x

42. Senkovich OA, Yin J, Ekshyyan V, Conant C, Traylor J, Adegboyega P, et al. Helicobacter pylori AlpA and AlpB bind host laminin and influence gastric inflammation in gerbils. Infect Immun. 2011;79(8):3106–16.10.1128/IAI.01275-10

43. Snelling WJ, Moran A, Ryan K, Scully P, McGourty K, Cooney J, et al. HorB (HP0127) is a Gastric Epithelial Cell Adhesin. Helicobacter. 2007;12(3):200–9.10.1111/j.1523-5378.2007.00499.x

44. Oleastro M, Menard A. The Role of Helicobacter pylori Outer Membrane Proteins in Adherence and Pathogenesis. Biology (Basel). 2013;2(3):1110–34.10.3390/biology2031110

45. Olson JW, Mehta NS, Maier RJ. Requirement of nickel metabolism proteins HypA and HypB for full activity of both hydrogenase and urease in Helicobacter pylori. Mol Microbiol. 2001;39(1):176–82.10.1046/j.1365-2958.2001.02244.x

46. Maier R, Benoit S. Role of Nickel in Microbial Pathogenesis. Inorganics.2019;7(7).10.3390/inorganics7070080

47. Ernst FD, Stoof J, Horrevoets WM, Kuipers EJ, Kusters JG, van Vliet AH. NikR mediates nickel-responsive transcriptional repression of the Helicobacter pylori outer membrane proteins FecA3 (HP1400) and FrpB4 (HP1512). Infect Immun. 2006;74(12):6821–8.10.1128/IAI.01196-06

48. Dressman J, Berardi R, Dermentzoglou L, Russell T, Schmaltz SB, J., Jarvenpaa K. Upper gastrointestinal (GI) pH in young, healthy men and women. Pharm Res. 1990;7(7):756–61.10.1023/a:1015827908309.

49. McConnell EL, Basit AW, Murdan S. Measurements of rat and mouse gastrointestinal pH, fluid and lymphoid tissue, and implications for in-vivo experiments. J Pharm Pharmacol. 2008;60(1):63–70.10.1211/jpp.60.1.0008

50. Hennigar SR, McClung JP. Nutritional Immunity: Starving Pathogens of Trace Minerals. Am J Lifestyle Med. 2016;10(3):170–3.10.1177/1559827616629117

51. Tsugawa H, Suzuki H, Matsuzaki J, Hirata K, Hibi T. FecA1, a bacterial iron transporter, determines the survival of Helicobacter pylori in the stomach. Free Radic Biol Med. 2012;52(6):1003–10.10.1016/j.freeradbiomed.2011.12.011

52. Waidner B, Greiner S, Odenbreit S, Kavermann H, Velayudhan J, Stahler F, et al. Essential role of ferritin Pfr in Helicobacter pylori iron metabolism and gastric colonization. Infect Immun. 2002;70(7):3923–9.10.1128/IAI.70.7.3923-3929.2002

53. Miles S, Piazuelo MB, Semino-Mora C, Washington MK, Dubois A, Peek RM, Jr., et al. Detailed in vivo analysis of the role of Helicobacter pylori Fur in colonization and disease. Infect Immun. 2010;78(7):3073–82.10.1128/IAI.00190-10

54. Timmerman MM, Woods JP. Potential role for extracellular glutathione-dependent ferric reductase in utilization of environmental and host ferric compounds by Histoplasma capsulatum. Infect Immun. 2001;69(12):7671–8.10.1128/IAI.69.12.7671-7678.2001

55. Zarnowski R, Cooper KG, Brunold LS, Calaycay J, Woods JP. Histoplasma capsulatum secreted gamma-glutamyltransferase reduces iron by generating an efficient ferric reductant. Mol Microbiol. 2008;70(2):352–68.10.1111/j.1365-2958.2008.06410.x

56. Kavermann H, Burns BP, Angermuller K, Odenbreit S, Fischer W, Melchers K, Haas R. Identification and characterization of Helicobacter pylori genes essential for gastric colonization. J Exp Med. 2003;197(7):813–22.10.1084/jem.20021531

57. Sun Y, Liu S, Li W, Shan Y, Li X, Lu X, et al. Proteomic Analysis of the Function of Sigma Factor σ54 in Helicobacter pylori Survival with Nutrition Deficiency Stress In Vitro. PLoS ONE. 2013;8(8).10.1371/journal.pone.0072920

58. Schweinitzer T, Mizote T, Ishikawa N, Dudnik A, Inatsu S, Schreiber S, et al. Functional characterization and mutagenesis of the proposed behavioral sensor TlpD of Helicobacter pylori. J Bacteriol. 2008;190(9):3244–55.10.1128/JB.01940-07

59. Eaton KA, Suerbaum S, Josenhans C, Krakowka S. Colonization of Gnotobiotic Piglets by Helicobacter pylori Deficient in Two Flagellin Genes. INFECTION AND IMMUNITY. 1996;64(7):2445–8.10.1128/iai.64.7.2445-2448.1996

60. Wu J-J, Sheu B-S, Huang A-H, Lin S-T, Yang H-B. Characterization of flgK gene and FlgK protein required for H pylori Colonization-from cloning to clinical relevance. World Journal of Gastroenterology. 2006 12(25):3989–399.10.3748/wjg.v12.i25.3989

61. Ottemann KM, Lowenthal AC. Helicobacter pylori uses motility for initial colonization and to attain robust infection. Infect Immun. 2002;70(4):1984–90.10.1128/IAI.70.4.1984-1990.2002

62. de Jonge R, Durrani Z, Rijpkema SG, Kuipers EJ, van Vliet AHM, Kusters JG. Role of the Helicobacter pylori outer-membrane proteins AlpA and AlpB in colonization of the guinea pig stomach. J Med Microbiol. 2004;53(Pt 5):375–9.10.1099/jmm.0.45551-0

63. Carlsohn E, Nystrom J, Bolin I, Nilsson CL, Svennerholm AM. HpaA is essential for Helicobacter pylori colonization in mice. Infect Immun. 2006;74(2):920–6.10.1128/IAI.74.2.920-926.2006

64. Koch MRA, Gong R, Friedrich V, Engelsberger V, Kretschmer L, Wanisch A, et al. CagA-specific Gastric CD8(+) Tissue-Resident T Cells Control Helicobacter pylori During the Early Infection Phase. Gastroenterology. 2023;164(4):550-

66. 10.1053/j.gastro.2022.12.016

65. Nguyen TL, Vieira-Silva S, Liston A, Raes J. How informative is the mouse for human gut microbiota research? Dis Model Mech. 2015;8(1):1–16.10.1242/dmm.017400

66. Xiang Z, Censini S, Bayeli PF, Telford JL, Figura N, Rappuoli R, Covacci A. Analysis of expression of CagA and VacA virulence factors in 43 strains of Helicobacter pylori reveals that clinical isolates can be divided into two major types and that CagA is not necessary for expression of the vacuolating cytotoxin. Infection and Immunity. 1995;63(1):94–8.10.1128/iai.63.1.94-98.1995

67. Arnold IC, Dehzad N, Reuter S, Martin H, Becher B, Taube C, Muller A. Helicobacter pylori infection prevents allergic asthma in mouse models through the induction of regulatory T cells. J Clin Invest. 2011;121(8):3088–93.10.1172/JCI45041

68. Kaebisch R, Mejias-Luque R, Prinz C, Gerhard M. Helicobacter pylori cytotoxin-associated gene A impairs human dendritic cell maturation and function through IL-10-mediated activation of STAT3. J Immunol. 2014;192(1):316–23.10.4049/jimmunol.1302476

69. Dailidiene D, Dailide G, Kersulyte D, Berg DE. Contraselectable streptomycin susceptibility determinant for genetic manipulation and analysis of Helicobacter pylori. Appl Environ Microbiol. 2006;72(9):5908–14.10.1128/AEM.01135-06

70. Cox J, Mann M. MaxQuant enables high peptide identification rates, individualized p.p.b.-range mass accuracies and proteome-wide protein quantification. Nat Biotechnol. 2008;26(12):1367–72.10.1038/nbt.1511

71. Tyanova S, Cox J. Perseus: A Bioinformatics Platform for Integrative Analysis of Proteomics Data in Cancer Research. In: v L. oS, editor. Methods in Molecular Biology: Humana Press; 2018.

72. Ferrero RL, Lee A. The Importance of Urease in Acid Protection for the Gastric-colonising BacteriaHelicobacter pyloriandHelicobacter felissp. nov. Microbial Ecology in Health and Disease. 2009;4(3):121–34.10.3109/08910609109140133

73. Skouloubris S, Labigne A, De Reuse H. Identification and characterization of an aliphatic amidase in Helicobacter pylori. Mol Microbiol. 1997;25(5):989–98.10.1111/j.1365-2958.1997.mmi536.x

74. Meister A, Tate S, Griffith O. Gamma-glutamyl transpeptidase. Methods Enzymol 1981;77:237–53.10.1016/s0076-6879(81)77032-0

75. Schmees C, Prinz C, Treptau T, Rad R, Hengst L, Voland P, et al. Inhibition of T-cell proliferation by Helicobacter pylori gamma-glutamyl transpeptidase. Gastroenterology. 2007;132(5):1820–33.10.1053/j.gastro.2007.02.031

76. Zhong Y, Anderl F, Kruse T, Schindele F, Jagusztyn-Krynicka EK, Fischer W, et al. Helicobacter pylori HP0231 Influences Bacterial Virulence and Is Essential for Gastric Colonization. PLoS One. 2016;11(5):e0154643.10.1371/journal.pone.0154643

77. Deutsch EW BN, Perez-Riverol Y, Sharma V, Carver J, Mendoza L, Kundu DJ, Wang S, Bandla C, Kamatchinathan S, Hewapathirana S, Pullman B, Wertz J, Sun Z, Kawano S, Okuda S, Watanabe Y, MacLean B, MacCoss M, Zhu Y, Ishihama Y, Vizcaíno JA The ProteomeXchange Consortium at 10 years: 2023 update. Nucleic Acids Res. 2023;51(D1):D1539–D48

78. Perez-Riverol Y BJ, Bandla C, Hewapathirana S, García-Seisdedos D, Kamatchinathan S, Kundu D, Prakash A, Frericks-Zipper A, Eisenacher M, Walzer M, Wang S, Brazma A, Vizcaíno JA The PRIDE database resources in 2022: A Hub for mass spectrometry-based proteomics evidences. Nucleic Acids Res. 2022;50(D1):D543–D52

